# Dissection of the role of a SH3 domain in the evolution of binding preference of paralogous proteins

**DOI:** 10.1101/2023.03.09.531510

**Authors:** Pascale Lemieux, David Bradley, Alexandre K Dubé, Ugo Dionne, Christian R Landry

**Affiliations:** Institut de Biologie Intégrative et des Systèmes (IBIS), 1030, avenue de la Médecine, Université Laval, Québec (Québec), Canada, G1V 0A6; Regroupement Québécois de Recherche sur la Fonction, l’Ingénierie et les Applications des Protéines, (PROTEO), 1045 Avenue de la Médecine, Université Laval, Québec (Québec), Canada, G1V 0A6; Centre de recherche en données massives (CRDM), 1065, avenue de la Médecine, Université Laval, Québec (Québec), Canada, G1V 0A6; Département de biochimie, microbiologie et bio-informatique, 1045 Avenue de la Médecine, Université Laval, Québec (Québec), Canada, G1V 0A6; Département de biologie, 1045 Avenue de la Médecine, Université Laval, Québec (Québec), Canada, G1V 0A6; Centre de Recherche du Centre Hospitalier Universitaire (CHU) de Québec-Université Laval, Québec, QC, Canada. Current address: Lunenfeld-Tanenbaum Research Institute, Sinai Health, Toronto, ON, Canada

**Keywords:** SRC Homology 3 (SH3) domain, gene duplication, ancestral sequence reconstruction, myosins

## Abstract

Protein-protein interactions (PPIs) drive many cellular processes. Some PPIs are directed by Src homology 3 (SH3) domains that bind proline-rich motifs on other proteins. The evolution of the binding specificity of SH3 domains is not completely understood, particularly following gene duplication. Paralogous genes accumulate mutations that can modify protein functions and, for SH3 domains, their binding preferences. Here, we examined how the binding of the SH3 domains of two paralogous yeast type I myosins, Myo3 and Myo5, evolved following duplication. We found that the paralogs have subtly different SH3-dependent interaction profiles. However, by swapping SH3 domains between the paralogs and by characterizing the SH3 domains freed from their protein context, we find that very few of the differences in interactions, if any, depend on the SH3 domains themselves. We used ancestral sequence reconstruction to resurrect the pre-duplication SH3 domains and examined, moving back in time, how the binding preference changed. Although the closest ancestor of the two domains had a very similar binding preference as the extant ones, older ancestral domains displayed a gradual loss of interaction with the modern interaction partners when inserted in the extant paralogs. Molecular docking and experimental characterization of the free ancestral domains showed that their affinity with the proline motifs is likely not the cause for this loss of binding. Taken together, our results suggest that the SH3 and its host protein could create intramolecular or allosteric interactions essential for the SH3-dependent PPIs, making domains not functionally equivalent even when they have the same binding specificity.

## Introduction

Protein domains are important structural and functional components of most proteins and often have modular and specific functions (Doolittle 1995). Among these domains are PDZ, SH2 and SH3 domains (Kuriyan and Cowburn 1997; Castagnoli et al. 2004) that direct protein-protein interactions (PPIs) by binding specific short linear motifs present on other proteins. SH3 domains (hereafter referred to as SH3s) have become powerful models to study the evolution of binding affinity and specificity because they are common and involved in many signaling pathways in eukaryotes (Kay et al. 2000; Dionne et al. 2022). The study of SH3 binding specificity and evolution is also relevant for biomedical applications. Indeed, SH3s bind short linear motifs and these motifs are often mutated in cancer and that pathogens use to take control of the signaling pathways of their host (Via et al. 2015; Dionne et al. 2022; Mihalič et al. 2023). Also, SH3s are found in several oncogenes and essential proteins for signal transduction (Latysheva et al. 2016; Massimino et al. 2021). Finally, SH3s can also be useful for biotechnological applications by enhancing binding between an enzyme and its substrate and for the assembly of synthetic protein scaffolds (Dueber et al. 2009; Park et al. 2022).

SH3s are numerous in the budding yeast (n = 27) and human proteomes (n > 300) (Xin et al. 2013; Teyra et al. 2017), largely as a result of the duplication of the genes that code for their host proteins (Vogel and Chothia 2006). Since the binding specificity of SH3s is key for the function of SH3-containing proteins, substantial effort has been put in place to map the determinants of their binding specificity. SH3 binding motifs most often correspond to PXXP (where X is any amino acid) and fall into canonical groups (Kuriyan and Cowburn 1997; Tonikian et al. 2009; Teyra et al. 2017). These groups sometimes overlap such that a given SH3 shares binding specificity with others. This, in principle, allows different SH3s to bind overlapping sets of proteins. However, it has been suggested that negative selection could act to limit spurious binding with non-cognate partners, which would prevent crosstalk between pathways with independent functions (Zarrinpar et al. 2003). Several other layers of regulation, including other interaction domains and protein colocalization, therefore combine with motif recognition to direct PPIs and allow a greater specificity (Dionne et al. 2022). This was recently illustrated by large-scale PPI binding efforts that have shown that SH3-directed PPIs *in vivo* also depend on the protein host and the position of the domain in the protein, which was defined as the protein context (Dionne et al. 2021). Since there are many mechanisms that contribute to the specificity of PPIs among SH3-containing proteins, one important question is how much do the protein-binding domains themselves contribute to the divergence of PPIs among paralogous proteins?

One powerful way to answer this question is to explore the evolution of the domains that have been preserved after their recent duplication. These domains and their host proteins had the same amino acid sequence at the moment of duplication, which makes it easier to understand how a relatively small number of amino acid changes contribute to a shift in specificity. Among the 27 SH3s present in 18 proteins of the budding yeast *Saccharomyces cerevisiae* are four pairs of paralogous domains that originated from a whole-genome duplication (WGD) event 100-150 million years ago (Wolfe and Shields 1997; Marcet-Houben and Gabaldón 2015; Wolfe 2015). Here, we focus on one of these protein pairs, the type I myosins Myo3 and Myo5.

Myo3 and Myo5 localize to actin cortical patches and are involved in endocytosis and exocytosis (Mochida et al. 2002). They both contain C-terminal SH3s and have overlapping sets of physical interactions in the Arp2/3 complex (Evangelista et al. 2000; Tarassov et al. 2008) and in actin polymerization networks (Lechler et al. 2000). From a genetics point of view, these two genes are at least partially redundant. The single deletion of either *MYO3* or *MYO5* has a slight effect on growth, but a double deletion leads to severe growth and actin cytoskeleton assembly defects (Goodson and Spudich 1995). Myo3 and Myo5 SH3s are a powerful model to study paralogous evolution because their multiple shared PPIs are critical for some cellular processes. The myosin paralogous SH3 domains have eleven different residues out of 59. These differences could have led to significant changes in the SH3 binding preferences because it has been found that a single substitution can lead to PPI disruption (Evangelista et al. 2000; Geli et al. 2000; Mochida et al. 2002; Dionne et al. 2021) In addition, they also have multiple known PPIs and a few specific PPIs, two for Myo3 and five for Myo5 (Szklarczyk et al. 2019), which could provide the resolution needed to dissect the divergence of specificity early after duplication.

In this study, we focus on the divergence of PPIs between Myo3 and Myo5 and examine the contribution of their SH3s to this divergence. We first examined the protein interaction network *in vivo* of the two paralogous proteins to identify their SH3-dependent PPIs. We then swapped the domains between the two paralogs to examine their contribution to PPI divergence. We also used resurrected ancestral SH3s to examine how the two domains have diverged since their duplication and pushed further in time to examine how they have evolved prior to the duplication event. Finally, to examine how the SH3s themselves may have changed, independently of the rest of the proteins, we examined the PPI profile of the SH3s *in silico* and free from their protein context.

## Results & Discussion

### SH3 domains play a major role in the function of Myo3 and Myo5 *in vivo*

The two paralogous genes *MYO3* and *MYO5* originated from the yeast WGD (Wolfe and Shields 1997). The corresponding amino acid sequences are 76% identical. They contain a N-terminal disordered region, a myosin motor, an ATP and actin-binding domain, and a C-terminal SH3 (Figure S7B, Myo3 : 1123-1181, Myo5 : 1088-1146). Eleven of the 59 residues of their SH3 are different (Figure S1D), which results in very similar structures as shown by the low root-mean-square deviation of atomic positions (RMSD = 0.386 Å, Figure S1B). RMDS < 3Å is only observed between closely related homologous structures (Chothia and Lesk 1986).

We first compared the PPI profiles of Myo3 and Myo5 *in vivo* in yeast. We used them as baits in a screen with a set of 296 proteins as preys. We measured pairwise PPIs using the dihydrofolate reductase protein-fragment complementation assay (DHFR PCA) (Tarassov et al. 2008) (Figure 1A). The preys were selected based on previously detected PPIs and the signaling network of either or both paralogs including 93 known interaction partners (see methods). Proteins that have been shown to interact ubiquitously in DHFR PCA are also included in the assay. We refer to them as abundance controls. These proteins create non-specific PPIs with most of the yeast proteins, likely through spontaneous DHFR fragment complementation. Thus, they can be used to detect changes in protein abundance (Tarassov et al. 2008; Levy et al. 2014) as PCA signal correlates with the amount of reconstituted DHFR enzyme (Freschi et al. 2013). Therefore, the more abundant a bait protein is, the stronger the PCA signal generated by the abundance controls will be. We included variants of Myo3 and Myo5, as baits, for which the SH3s were encoded in codon-optimized sequences (optSH3s) for expression in *S. cerevisiae* since we also test codon-optimized ancestral SH3s below. Also, because our goal was to examine the role of the SH3s in the divergence of PPIs, the experiment included the SH3-depleted paralog variants for which a short flexible peptide (GGSSGGGG) was used to replace the SH3s. We used these constructs as baits to determine which PPIs are SH3-dependent (Figure 1A) (Dionne et al. 2021).

**Figure 1.**
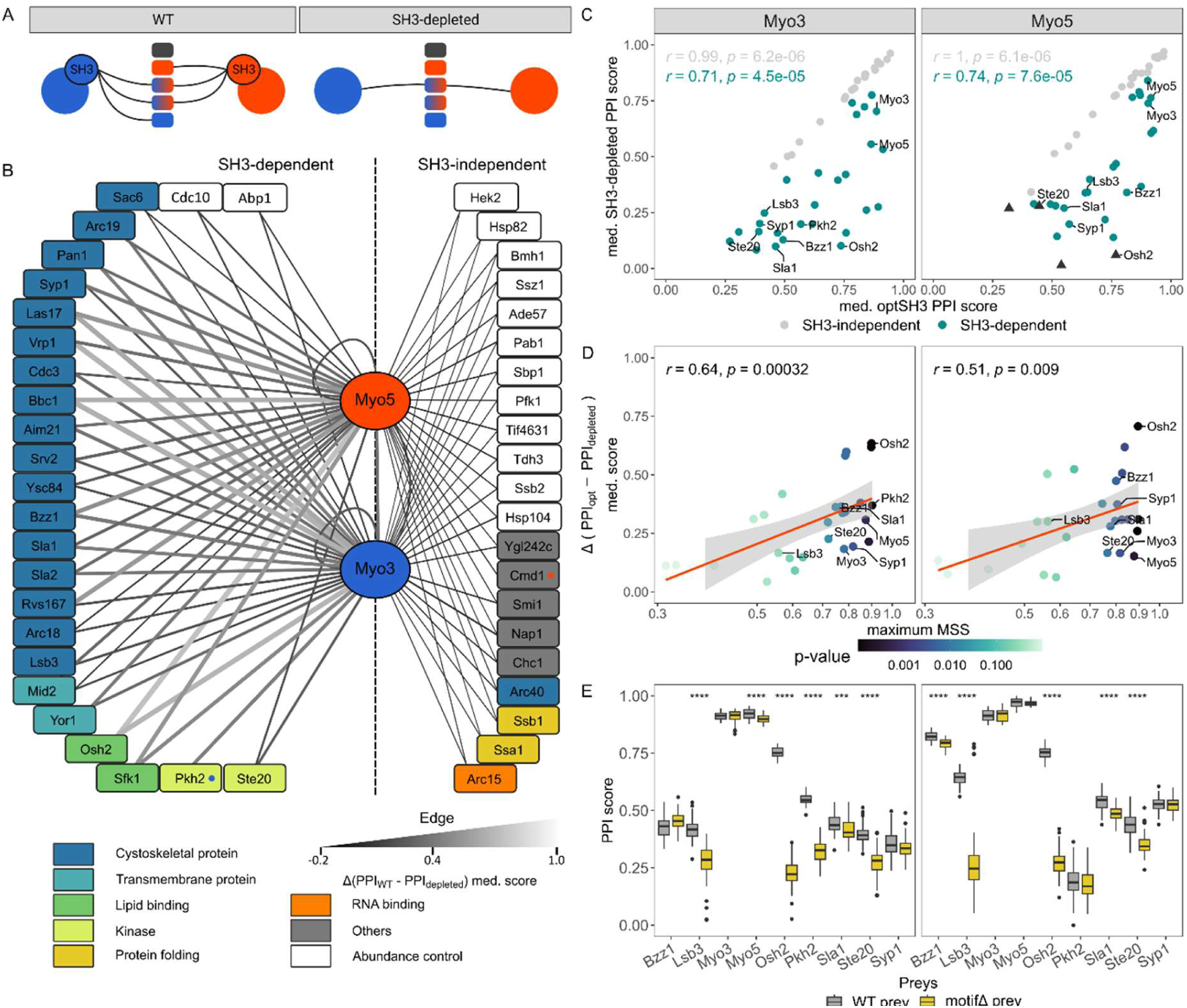
Characterization of the interactome of Myo3 and Myo5. **A.** Schematic representation of the experimental design. The baits are shown in circular shape and preys are shown in rectangle shape. PPIs are represented by the lines and the color of the prey nodes shows the binding preference of each paralog. The left panel represents the WT PPI network of the paralogs and the right panel shows the PPI network of the SH3-depleted paralog variants. The comparison between both experiments allows us to identify SH3-dependent interactions. **B.** Myo3 and Myo5 PPI network. The bait proteins are displayed in the center of the network. The edges show the SH3 contribution to the PPI: Δ(PPI_WT_ - PPI_depleted_) median score. The vertical dash line separates the SH3-dependent, for one or both paralog, and SH3-independent preys as determined by Wilcoxon tests. Paralog specific partners are indicated by a colored dot in the nodes (Myo3 : blue, Myo5 : red). Node colors indicate functions (Cherry et al. 2012). White nodes are abundance controls. **C.** Scatter plots comparing the PPI scores of the optSH3 paralog variants with the SH3-depleted paralogs. Myo3 data are shown on the left and Myo5 data are shown on the right as it is for panel D and E. Each data point represents the median score (8 or 10 biological replicates) of an interaction with an individual partner. Triangle-shaped data points are PPIs with fewer replicates (n ≤ 2) which prevented us from concluding if they were SH3-dependent. The labeled data points are preys that were used to confirm the proline motif predictions in panel E. **D.** Scatter plot comparing the Δ(PPI_opt_ - PPI_depleted_) median score with the corresponding motif prediction score of the preys (maximum MSS). The color scale represents the p-value of the motif predictions. **E.** PPI scores (48 biological replicates) of the WT preys and motifΔ preys with the extant paralogs. Wilcoxon tests were performed between the extant and the motifΔ preys and significance levels are show for each significant comparison (*** : p <= 0.001, **** : p <= 0.0001).

We detected most of the previously reported PPIs in three different PPIs databases (83/135) (Figure S2A) (Stark et al. 2006; Licata et al. 2012; Szklarczyk et al. 2019) and the majority of those detected by more than one method (78/95) using the BioGRID as a reference (Stark et al. 2006). Excluding the abundance controls for which information is absent from the reference dataset, only one interaction was detected for the first time (Figure S2B). Myo3 and Myo5 share most of their interaction partners (Figure 1B). Excluding the abundance controls, the paralogs share 32 PPIs and they have one specific PPI each (Myo3 : Pkh2, Myo5 : Cmd1, Figure 1B, Figure S4A). Moreover, we observe significant differences in PPI strength for 13 of the shared ones. However, specific PPIs identified in previous work were not detected in our assay. Broadly, PPI mapping thus confirms the common function of the paralogs in the actin cytoskeleton organization (Goodson and Spudich 1995; Lechler et al. 2000) and their functional redundancy. Also, several PPIs and the quantitative nature of the PCA signal were validated by performing small-scale validation experiments in individual liquid cultures (Spearman, optSH3s r = 0.92, p = 3.3×10-6, Figure S3F).

The comparison between the median interaction scores (PPI score, see method) of the WT codon and the optSH3 reveals very strong correlations for both paralog (Spearman, r ≥ 0.99, p ≤ 1.6×10^-6^) and the optSH3s and WT SH3s PPI scores have very similar medians (WT Myo3 : 0.764, optMyo3 : 0.756, WT Myo5 : 0.765, optMyo5 : 0.769) suggesting that the codon optimization has no effect on the DHFR PCA results. Thus, the data obtained with the optSH3 paralog variants were used as reference to analyze further experiments testing codon-optimized SH3s. Henceforth, we refer to the WT domains as extant domains (extantSH3s) as we also discuss ancestral domains. We identified SH3-dependent partners by analyzing DHFR PCA signals of the optSH3 paralog variants compared to the SH3-depleted variants (Figure 1C). PPIs were classified as SH3-dependent when a significant difference in PPI scores was observed between the SH3-depleted and optSH3 paralog variants (Wilcoxon, Benjamini-Hochberg corrected p-value < 0.05). We observed for Myo3, 25 SH3-dependent PPIs and 21 for Myo5. Excluding the Myo3-specific PPI with Pkh2, the three additional SH3-dependent PPIs of Myo3 can be explained by the lack of PCA signal for the SH3-depleted Myo5 variant, which prevents the statistical test from being conclusive (Figure S4B). Also, the paralogs interact with each other, displaying a modest but significant SH3-dependent property. Some of these SH3 dependencies support results from previous studies. For example, as we observe here when the entire domain is removed, a previously characterized mutation in the SH3s disrupts SH3-dependent PPIs (W1158S) between Myo3 and Las17 (Evangelista et al. 2000) and Myo5 (W1123S) with Bbc1 and Vrp1 (Geli et al. 2000; Mochida et al. 2002)(Figure 1B).

We manually reviewed the function of each interaction partner using the *Saccharomyces* Genome Database (SGD) (Cherry et al. 2012). This analysis revealed that the SH3-dependent partners are mostly involved in the cytoskeleton assembly whereas the SH3-independent partners are involved in broader cellular functions or are the abundance controls (Figure 1B). More than half (14/23) of the SH3-independent PPIs involve the abundance controls, which are not expected to be dependent on the bait protein, leaving only nine specific SH3-independent PPIs. Also, we have to mention that some SH3-independent PPIs partners could interact ubiquitously. For example, the abundance control Ssb2 and its paralog, Ssb1, are known to have a similar function as cytosolic chaperons, but Ssb1 is not classified as an abundance control (Werner-Washburne et al. 1987; Nelson et al. 1992) (Figure 1B). The finding that many of the Myo3 and Myo5 PPIs depend on their SH3s confirms that the SH3s have an important role in the function of these two proteins (Geli et al. 2000).

To validate our observations on SH3 dependency, the putative proline-rich binding motifs were predicted on the SH3-dependent interaction partners. Predictions were done using a specificity matrix of the preferred binding peptides determined by peptide phage display (Tonikian et al. 2009). A motif prediction score (maximum Motif Similarity Score (MSS)) between 0 and 1 was obtained for each prey by identifying the protein sub-sequence with the strongest match to the Myo3/Myo5 specificity matrix. A positive correlation was observed between the SH3 contribution to the interaction, i.e. the difference of PCA signal between the optSH3 and SH3-depleted protein variants (Δ(PPI_opt_ - PPI_SH3-depleted_) median score), and the motif prediction score for both paralogs (Spearman, Myo3 r = 0.64 p = 3.2×10^-4^, Myo5 r = 0.51 p = 9.0×10^-3^, Figure 1D). This supports our prediction that SH3-dependent PPIs depend on the match of the cognate Myo3 and Myo5 SH3 binding motifs to their interaction partners, although other major factors such as prey abundance could contribute to variation in PCA signal. To further validate the SH3 and motif dependency of PPIs, we used the same linker sequences used above to delete the SH3s but this time to replace the motifs with the highest MSS score on interaction partners, creating motifΔ preys. We successfully modified nine of these motifs and measured their impact on PPIs by DHFR PCA (Table S5, S11 in File S1).

A significant decrease in interaction strength for six partners of Myo3 and five for Myo5 was observed when the top-scoring proline motif was replaced with the linker. SH3-dependent PPIs are therefore most likely dependent on the top-scoring proline motif on the interaction partners (Figure 1E). Cases where no dependency or partial loss of PPI is observed, could be due to redundancy in the poly-proline motifs as many of the interaction partners contain multiple proline-rich motifs (Figure S7B), or to the contribution of the disordered regions surrounding the motif to the PPI. These regions are known to be involved in PPIs and to favor multiple binding conformations, as we can hypothesize for Myo3 and Myo5 (Kelil et al. 2016; Morris et al. 2021). For example, we detected a motif dependency for the interaction of Myo3 with the Myo5 motif but not the reciprocal (Figure 1E). However, there are many proline motifs in the disordered regions in Myo3 and in Myo5 (Figure S7B). One possibility is that the replacement of the proline motif in Myo3 was masqued by one of the other proline-rich motifs in proximity, but it would not be the case for the PPI of Myo3-Δmotif with Myo5. Indeed, Myo5 has five alternative motifs and Myo3 has only three alternative motifs that could explain the differences in motif dependency (Figure S7B). Furthermore, the mechanisms behind the contribution of disordered regions to PPI are also undefined, but the modifications of these regions, as we did by replacing the binding motif, can also affect binding (Bugge et al. 2020).

### SH3-dependent interaction differences between paralogs are not explained by the SH3 contribution to the PPIs

We compared the Myo3 and Myo5 interaction datasets and we observed that Myo5 raw PCA signal is stronger on average than Myo3 raw PCA signal (Figure S3B, Wilcoxon p < 2.2×10^-16^). Since DHFR PCA measures the amount of protein complex formed (Remy and Michnick 2007; Freschi et al. 2013), differences in binding could in principle result from differences in bait abundance. In order to confirm that this bias is due to protein abundance, we measured protein abundance using Myo3 and Myo5 GFP-tagged strains (Figure 2A). The results reveal that Myo5 is slightly more abundant than Myo3 (Figure 2A, Student’s t-test, p = 6.1×10^-4^), as reported in public databases (Cherry et al. 2012; Breker et al. 2013). We also included the SH3-depleted proteins to examine if the deletion of the SH3s affected abundance. Dionne et al. showed that SH3 depletion had minimal to no effect on abundance by Western blotting for many SH3-containing proteins. We observe the same results here using flow cytometry (Figure 2A, Student’s t-test, Myo3 p = 0.92, Myo5 p = 0.46). Thus, to compare the interaction strength of the two paralogs, we had to ensure that the normalization method to compute the PPI score eliminated the effect of bait abundance on the interaction data (see method). Indeed, the median PPI scores of the abundance controls with the paralog variants confirm that the abundance bias was removed by the normalization (Figure S3C, Wilcoxon p = 0.96).

**Figure 2.**
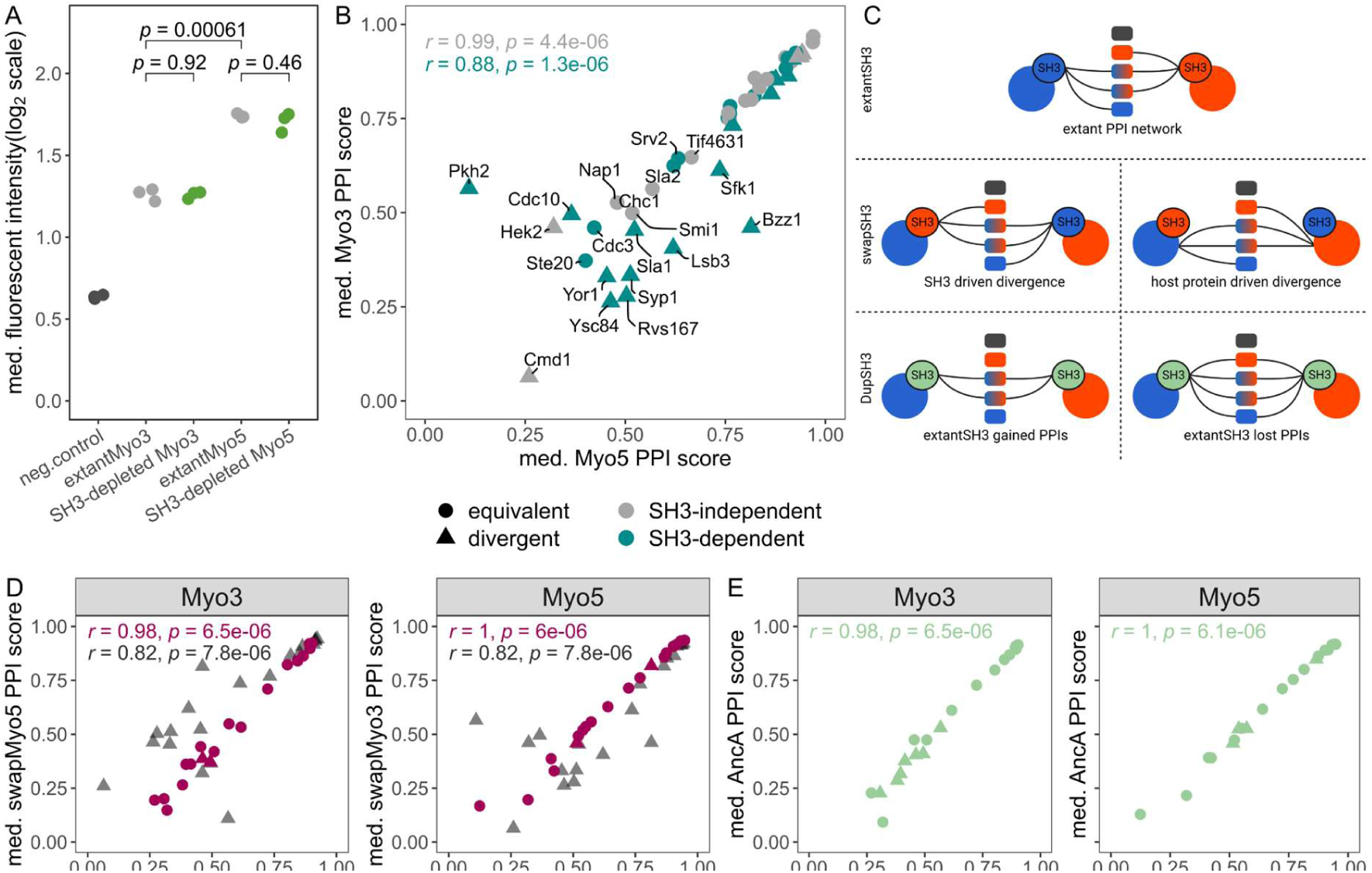
SH3 domains are not sufficient to explain the PPI preferences of the paralogs. **A.** Expression level of extant paralogs and SH3-depleted protein variants tagged with GFP. The median cell fluorescence signal (arbitrary units) is measured for 5000 individual cells from three independent cultures (individual data point). The negative control is a strain not expressing GFP (strain BY4741). Student’s t-test p-values are shown above each comparison. **B.** Comparison of the optSH3 variants interaction profiles of Myo3 and Myo5. The colors represent the SH3 dependency of the PPIs. Triangle-shape marks are divergent PPIs between the paralogs. **C.** Schematic representation of the expected PPI network of the swapSH3 and AncA paralog variants compared to the extant PPI network under different evolutionary scenarios. The PPIs are represented by lines and the colors of the nodes show the extant paralog partners. The swapSH3 determines which paralog-specific PPI can be recovered only by the contribution of its native SH3. The PPIs observed with AncA identifies which paralog-specific PPI was gained or lost following the duplication event. **D and E.** Comparison of the PPIs of Myo3 and Myo5 containing various SH3s. The host protein is shown at the top of each plot. **D** shows the comparisons between the swapSH3 (y-axis) and the optSH3 (x-axis) in fuschia. The grey data points represent what we would expect if the divergence was SH3 driven, which would allow the swapSH3 to recover all the extant SH3-dependent PPIs and replicate the PPI profiles of the SH3 in their respective paralog context. **E** shows the comparisons of the AncA (y-axis) with optMyo3 or optMyo5 (x-axis) in both paralogs. Spearman correlation coefficients are shown on the plots.

We quantified the similarity in binding profiles of Myo3 and Myo5 by directly comparing the normalized PPI scores of the two extant proteins. This comparison reveals a strong correlation for SH3-dependent interaction scores (Spearman, r = 0.91, p = 1.4×10^-6^) and an even stronger correlation for SH3-independent ones (Spearman, r = 0.99, p = 4.4×10^-6^) (Figure 2B). However, there remain a few important differences among the SH3-dependent PPIs. For example, Bzz1 is a strong binding partner for Myo5 but a weak one for Myo3, and we observe the opposite pattern for Pkh2 (Figure 2B). Most of the differences between the two paralogs are observed for PPIs with weaker signals. DHFR PCA signals can reach saturation for stronger PPIs (Remy and Michnick 2007), potentially preventing us from detecting divergence for the high-intensity binding partners. Furthermore, if the divergence in SH3s is contributing to the divergence of binding, we would expect those to be for cases of weak PPIs as SH3s form weak transient PPIs with specific proline motifs (Dalgarno et al. 1997; Kay et al. 2000). Sequence divergence in the SH3s could thus affect these PPIs most strongly. These results suggest that the difference in binding between Myo3 and Myo5 could potentially be driven by changes in their SH3.

In order to determine whether the SH3s are sufficient or not to confer PPI specificity to the paralogs, we swapped the SH3s between them (Figure 2C). A paralogous SH3 inserted in the non-native paralog is referred to as a swapSH3 paralog variant. If the SH3s are sufficient to confer PPI specificity, swapping the domains will also switch the SH3-dependent PPI profile of the paralogs. Any differences caused by the paralogous SH3s could result from sequence changes following their last common ancestor resulting in gains or loss of interaction strength of either paralog (Figure 2C). To differentiate gains and losses of PPIs, the ancestral domain at the last common ancestor node (AncA) was resurrected and its codon-optimized sequence was also inserted in both paralogs. The ancestral sequence reconstruction (ASR) was performed using a maximum likelihood-generated phylogeny across fungal Myo3/Myo5 orthologs with Fast-ML (See methods) (Pupko et al. 2000; Ashkenazy et al. 2012) (Figure S1A, Data S2). We included only fungal orthologs in the phylogeny because sequence similarity drops as low as 39.3% between orthologs in this group. Adding more divergent orthologs from outside the fungal kingdom would have added little further insight into the Myo3/Myo5 evolutionary history post duplication. The Myo3/Myo5 tree topology is supported by strong phylogenetic branch supports (aLRT), and all ancestral sequences tested were reconstructed with high posterior probability (Figure S1C). Myo3 and Myo5 SH3 orthologous sequences are mostly conserved differing on 11/59 positions, but we observe that 20/59 positions diverged in at least one ortholog since their last common ancestor (Figure S1D). Therefore, the AncA has high sequence identity with both extant SH3s (53/59 Myo3, 54/59 Myo5, Figure S1E). The AncA sequence was reconstructed with high confidence (aLTR = 96.6) and 58/59 positions are predicted with 95% or higher posterior probability (Figure S1C). The insertion of swapSH3 and of AncA in the extant paralogous contexts create chimeras that will generate PPIs with the proteins present in the cell. These artificial PPIs will allow us to investigate the limits of the SH3 domain sequence in the binding of its interaction partners.

Swapping the SH3s and inserting AncA in Myo3 and Myo5 reveal that the interaction profiles of the two paralogs are not affected by the difference in the sequence of their paralogous SH3s and their ancestor (Figure 2D, 2E). The paralogs with swapped SH3s have very similar SH3-dependent interaction profiles as their native ones (Figure 2D, Spearman, Myo3 Myo3 r = 0.98 p = 6.5×10^-6^, Myo5 r = 1 p = 6×10^-6^). Previously, we have found that the PPI intensity between paralogs is different for 13 cases out of 25 SH3-dependent PPIs (Figure S2A). For instance, Pkh2 interacts with Myo3 but not with Myo5, at least with our assays. This specific PPI with Myo3 is SH3-dependent, indicating that the SH3 plays an important role in this PPI. Thus, we expected to detect a similar PPI intensity with Pkh2 when the Myo3 SH3 was inserted in Myo5, but we observed no gain in PPI with the Myo5 swapSH3 variant (Figure S2C). In fact, it was previously found that the protein context affects the PPI profiles of SH3 domains (Dionne et al. 2021). Yet, in some cases, swapping the SH3s between two proteins induced novel specific interactions to the new host protein (Dionne et al. 2021). Thus, we expected that the effect of the protein context would be reduced with closely related proteins, but, despite the SH3 weight in the Pkh2 PPI and the high sequence similarity between the paralogs, the swapSH3 is not able to transfer its contribution to create a Myo5-Pkh2 PPI. The same result is obtained with the insertion of AncA in both paralogs (Figure 2E, Spearman, Myo3 r = 0.98 p = 6.5×10^-6^, Myo5 r = 1 p = 6.1×10^-6^). Overall, these results show that, although different in sequence, the two paralogous domains and their last common ancestor SH3 do not contribute to differences in binding between the two paralogous proteins Myo3 and Myo5.

### Ancestral SH3 domains progressively lose SH3-dependent PPIs

Our results comparing Myo3 and Myo5 PPIs showed that they have quantitatively slightly different interaction profiles, especially for SH3-dependent PPIs (Figure 2B). However, none of the differences observed appears to be driven by the divergence of the domains themselves. Surprised by this, we wondered how robust PPIs of these two paralogs were to the identity of their SH3s. We therefore resurrected more ancestral sequences at speciation nodes (AncB, AncC, AncD) that predate the last common ancestor of Myo3 and Myo5 and examined PPIs again (Figure 3A). AncB is the SH3 that was present in the last common ancestor of all the WGD species. Some of these species lost the second copy of the Myo3/Myo5 orthologs. AncC and AncD are, respectively, the last common ancestors for the *Saccharomycetaceae* (family) and *Saccharomycetales* (order) clades. The same ancestral sequence reconstruction method as the one used for AncA was applied to infer AncB, AncC and AncD domains. High aLRT and strong topology support was obtained even for the oldest node used for the ancestral reconstruction (aLTR, AncB = 85.6, AncC = 89, AncD = 99.9, Figure S1A). However, AncD’s posterior probability distribution is slightly lower than the more recent ancestral SH3s, which is expected due to the higher sequence diversity used for the reconstruction (Figure S1C, Data S2). Sequence identity gradually decreases to 55% between the extant Myo3 SH3 and AncD, and to 50% for Myo5 SH3 compared to AncD (Figure S1E). To support the ASR predictions, we used AlphaFold2 to predict the ancestral SH3 3D structures (Figure 3B, Data S3) (Jumper et al. 2021). The predicted structures show very low RMSD when compared to the extantSH3s, indicating overall structure conservation despite the low sequence identity.

**Figure 3.**
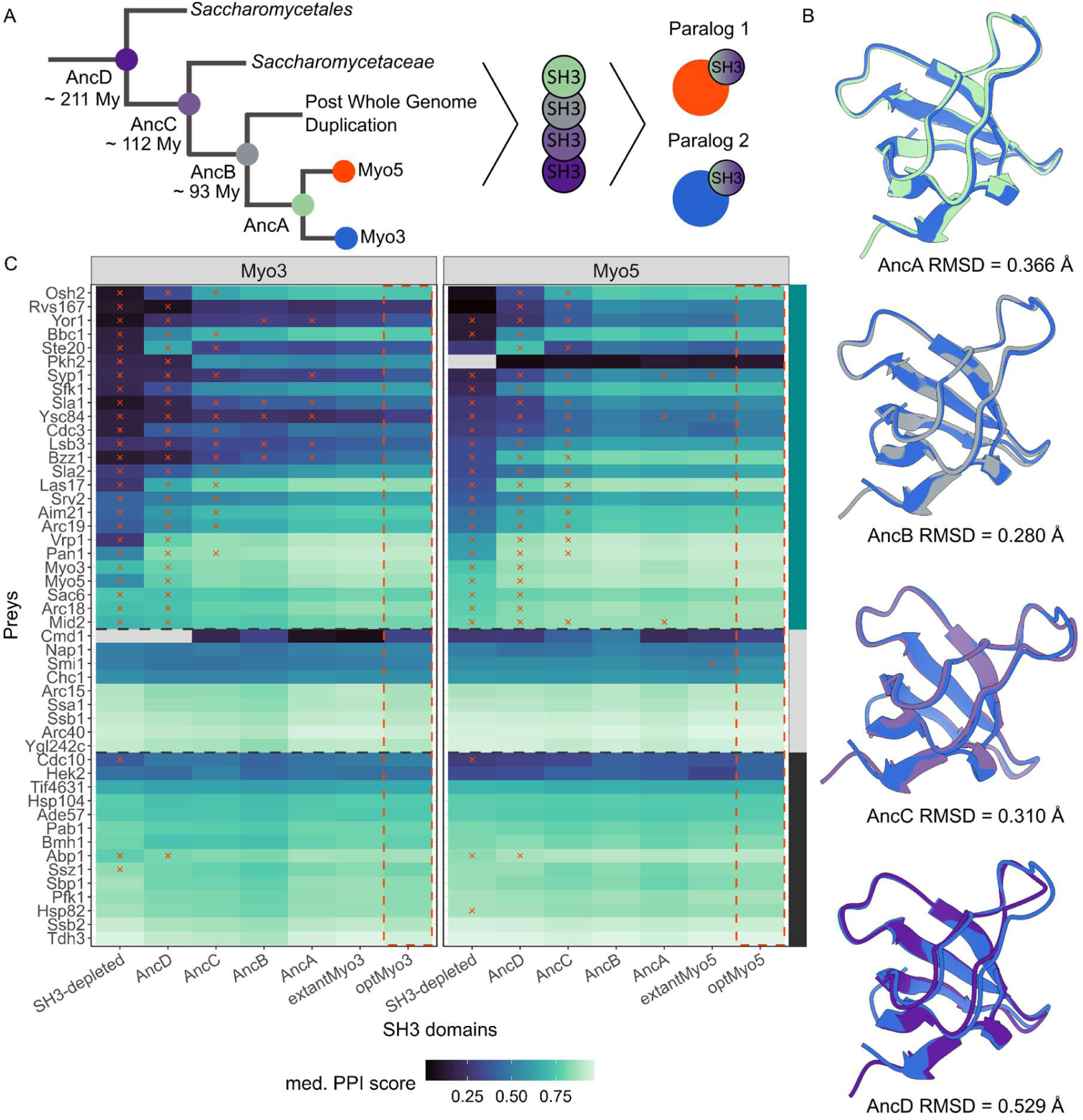
Ancestral SH3 domains affect the PPIs of Myo3 and Myo5. **A.** Simplified phylogenetic relationship of yeast where nodes represent an extant SH3 (blue and red) or an ancestral SH3 that was reconstructed with an estimation of the time at which these domains existed. **B.** Superimposed extantMyo3 SH3 structure (blue, PDB:1RUW) and the AlphaFold2 predicted structures of each reconstructed ancestral SH3 showing high structural conservation as estimated by RMSD. **C.** Heatmap showing the PPIs of SH3 variants inserted in Myo3 and Myo5 with each of their interacting partners (preys, y-axis). The preys are ordered by SH3-depleted Myo5 PCA signal and by interaction type with the paralogs (right annotation, cyan : SH3-dependent for one or both paralogs, gray : SH3-independent, black : abundance controls). The tile color corresponds to the median PPI score and the red x marks correspond to a significant difference (p < 0.05, Wilcoxon test, Benjamini-Hochberg correction) in interaction strength compared to the control interaction (dashed red frame). Missing data are shown in gray.

By measuring PPIs of Myo3 and Myo5 containing these ancestral sequences, we observe a general and progressive loss of signal as domains get more ancient, specifically for SH3-dependent PPIs (Figure 3C). For the most ancient SH3, the PPI network shows a significant interaction loss for its SH3-dependent PPIs compared to the optSH3s (paired-Wilcoxon, Myo3 p = 5.5×10^-6^, Myo5 p = 2.4×10^-7^). However, no difference is observed for the SH3-independent median PPI scores across SH3-depleted, AncD and optSH3 variants (Kruskal-Wallis, Myo3 p = 0.47, Myo5 p = 0.98). The pattern of loss for AncD PPI indicates that the ancestral domain loses PPIs in a similar way to the SH3-depleted paralogs (Figure 3C). Since SH3-independent PPIs with the abundance controls appear to be preserved, the gradual loss of interaction is unlikely to be caused by protein instability and degradation. To validate this hypothesis, we tagged the paralog variants with GFP and measured protein abundance. Protein abundance was not significantly different compared to the extant paralogs for any of the variants (Figure S3E). Therefore, the results point towards an actual loss of binding by the ancestral domains. An exception in the general tendency is observed for the prey Ste20, which gains interaction with the AncD paralog variants (see below).

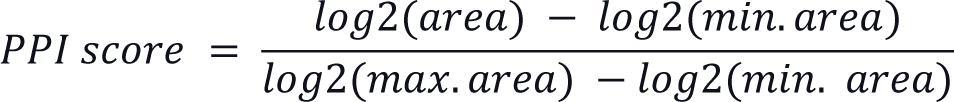

Since the SH3s are surrounded by disordered regions containing proline motifs, we explored the hypothesis of intramolecular binding mediated by the ancestral SH3s to explain the loss of PPI with the extant partners. It was shown that AlphaFold2 is able to predict intramolecular binding of a peptide chain containing a SH3 followed by a disordered linker and a proline motif (Tsaban et al. 2022). Thus, we used AlphaFold2 to predict the structure of the paralogs carrying the ancestral SH3s (AncA to AncD, DataS3) (Jumper et al. 2021). These analyses did not predict intramolecular binding. In addition, we performed AlphaFold Multimer with Myo3, Myo5 and AncD SH3 domains and their surrounding regions in the paralogs to assess if another proline motif could be favored for the binding of AncD variants (Figure S7B, Myo3 : 961-1272, Myo5 : 961-1219) (Evans et al. 2022). However, AncD is predicted to bind the same motifs as the extant SH3s (DataS3). Thus, despite the predictions, we are not able to define the role of the disordered regions in the binding of the SH3s. It is known that AlphaFold2 predictions are less reliable for the prediction of disordered regions because the algorithm considers coils as unstructured, which can lead to overestimation of disordered regions (Wilson et al. 2022). In our case, this bias in the prediction could prevent us from detecting a transient intramolecular or intermolecular binding between a proline motif and the ancestral

We performed an orthology analysis on the SH3-dependent interaction partners of the myosins to examine if binding to the same partners as the extant duplicates could have taken place in the clade defined by AncD. We found that most of the interaction partners have orthologs in the species included in AncD clade (Figure S5A). Thus, AncD interaction loss with the extant PPI partners can not be explained by the absence of their ancestors at AncD time of existence. We also verified the conservation of the *in vivo* validated proline motifs (Figure 1E) on PPI partner orthologs (Figure S5B). We found that proline motifs are present on the PPI partners’ orthologs in AncD clade, except for Pkh2. The case of Pkh2 is interesting because we showed that it was a Myo3 specific interaction partner. Pkh2 has a paralog, Pkh1, which was included in the DHFR PCA experiment, but showed no PPI with either myosins. Since Pkh2 binds specifically to Myo3, we could speculate that this is a SH3-dependent interaction gain of Myo3 since the WGD event. This hypothesis is also supported by the motif conservation analysis, because very few proline binding motifs are predicted on the orthologs of Pkh2 outside of the WGD clade, indicating that it was not interacting with the ancestral myosin (Figure S5B). Overall, the interaction partners orthology analysis and the motif conservation analysis suggest that AncD was binding to the most extant interaction partners’ ancestors.

Furthermore, we investigated the surface properties of the SH3 domains to examine the possibility that the surface outside of the SH3 binding site could contribute to a transient intramolecular conformation with the rest of the paralogs. We compared hydrophobicity and electrostatic potential of Myo5 SH3 and AncD surfaces using ChimeraX metrics (Figure S6) (Laguerre et al. 1997; Pettersen et al. 2021). The same analyses were performed using Myo3 SH3 and very similar results to Myo5 SH3 were obtained due to the high sequence similarity between the two extant SH3s. Thus, only Myo5 SH3 and AncD surface properties are illustrated in Figure S6. Hydrophobicity properties are conserved between the two SH3s. However, we observed that a surface patch with negative electrostatic potential on Myo5 SH3 is neutral on AncD (Figure S6). This surface change could potentially affect intramolecular binding and explain our results but this would require further investigation. We also performed an intramolecular coevolution analysis on the full-length proteins using EVcouplings (v0.2, (Hopf et al. 2019)), but no significant coevolution signal was detected between the SH3 domain and the rest of the myosins (Data S7). In short, multiple *in silico* predictions were performed, but none of the approaches allowed us to discern the mechanism behind AncD loss of binding.

### SH3 variants affinity to proline motifs does not explain the divergence between the paralogous PPIs

The general decrease of binding for ancestral sequences, particularly of AncD, could be caused by a diminished binding affinity of this ancestral domain with the extant binding motifs. Using the binding motif prediction on SH3-dependent preys validated above (Figure 1D, E), we performed *in silico* SH3-peptide docking with the ancestral and extant SH3s to test this hypothesis.

The experimental structures of the extant Myo3 (PDB:1RUW) and Myo5 (PDB:1YP5) SH3s and the predicted AncD structure (Figure 3B, Data S3) were used for computational molecular docking (Geng et al. 2017) with peptides corresponding to binding motifs (10 amino acids long) from the preys (Table S5 in File S1). Nine of these binding motif predictions were used to validate the SH3 dependency (Figure 1D, E). Multiple structures were examined each time and they were sorted into clusters based on RMSD. The clusters and structures were ranked by the docking scoring algorithm (Geng et al. 2017). Next, a global energy minimization step was applied to the best structures of each cluster and the SH3-peptide interaction energies were computed (ΔG (kcal/mol)) (Schymkowitz et al. 2005). In 43% of the docking peptide-SH3 combinations, the best-ranked cluster included the lowest ΔG structures (Myo3: 8/28, Myo5: 15/28, AncD: 13/28). Additionally, we observed various conformations in the docked-peptide structures even if these structures are grouped in the same cluster (Figure 4A). We thus computed the median ΔG of the best structures of each cluster to consider this variability (see methods). Using the median ΔG to compare the clusters, we validated that the ranking of the clusters is a good indicator of the binding energy. Indeed, the first rank cluster shows a lower distribution of median ΔG compared to clusters with a higher rank for each SH3 (Figure S8B). Thus, we continued the analysis with the median ΔG of the top cluster of each docking as a proxy for the affinity.

**Figure 4.**
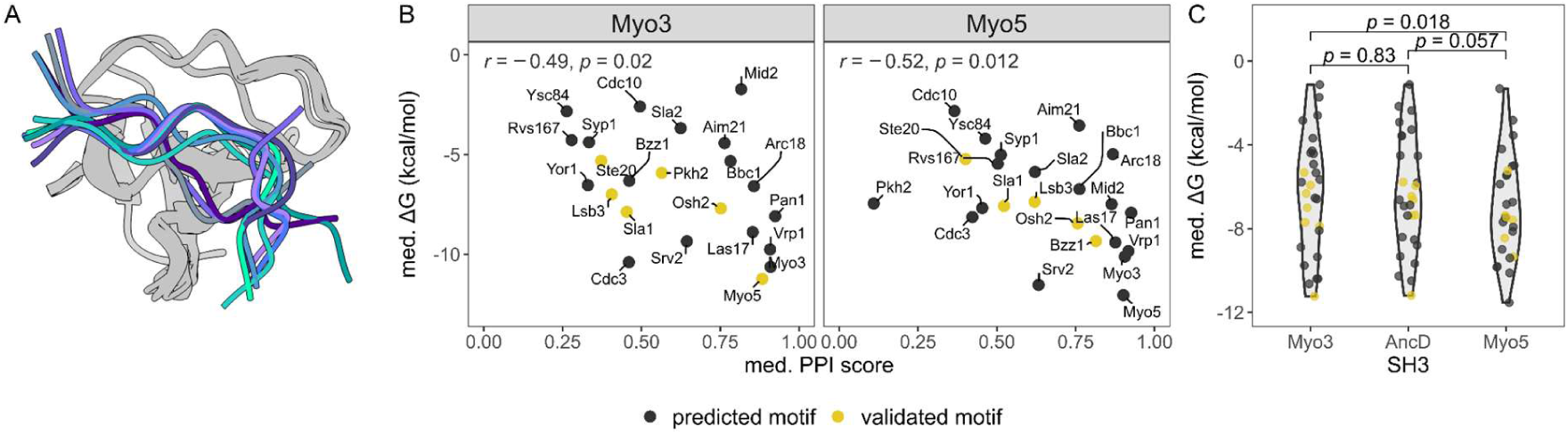
Contribution of SH3 binding to their motifs estimated by molecular docking. **A.** Superimposed structures of the ten best-scored docking structures of Osh2 binding peptide on Myo3 SH3. The peptides show a variety of binding conformations whereas the SH3 structures remain similar. **B.** Scatter plots showing the relationship between the computed median ΔGs of the docked peptide-SH3 complex and the corresponding PPI scores of the extant paralogs as measured by DHFR PCA. Yellow data points correspond to proteins for which the proline motifs have been validated *in vivo* (Figure 1E). Spearman correlation coefficients (r) and p-values appear at the top of each plot. **C**. Violin plot comparing the distribution of median ΔG values for each SH3s. Paired-Wilcoxon tests p-values are labeled on top of the violin plot.

We observe significant correlations between the median ΔGs and the PPI scores obtained by DHFR PCA for both paralogs (Spearman, Myo3 r = -0.49 p = 0.02, Myo5 r = -0.52 p = 0.012) (Figure 4B). This result supports the binding motif predictions illustrated in Figure 1D given that the peptides used for structural modeling derive from our binding motif predictions. To examine if the paralogous SH3s have different binding affinities to the proline motifs, we compared the median ΔGs between Myo3 and Myo5 SH3s (Figure 4C). This comparison suggests that Myo5 SH3 binds globally more strongly to the proline motifs (predicted and validated) than Myo3 SH3 (paired-Wilcoxon, p = 0.018). In addition, the comparisons between AncD and the extantSH3s median ΔGs do not explain the data collected *in vivo.* Indeed, there is no significant decrease in binding affinity computed for AncD compared to the extantSH3s (paired-Wilcoxon, Myo3 p = 0.83, Myo5 p = 0.057) while a general decrease in binding is detected *in vivo* for the AncD-paralogs variants. We note however that in these comparisons, AncD appears to have median ΔGs more similar to that of Myo3 than of Myo5 SH3s, although the differences are not significant. Thus, even if the docking reveals a globally stronger binding of the proline motifs to Myo5 SH3 compared to Myo3 SH3, it does not explain the differences in the binding profile of the extant SH3s *in vivo*. The inconclusive molecular docking results and the fact that they are not in agreement with the *in vivo* experiments led us to seek validation with additional *in vivo* experiments.

The conflicting results between the PCA experiment and the molecular docking do not allow us to conclude as to whether the SH3 affinity to the proline motifs is one of the main factors shaping the differences in PPIs of Myo3 and Myo5, nor to explain the loss of interaction shown with the replacement of native SH3 by the AncD variant in the paralogs. Another hypothesis that could explain the PCA results is that the PPIs of the ancestral SH3 with the preys are reduced because a SH3 also interacts (intramolecular interaction or allosterically) with its host protein in a way that decreases its ability to bind. The context in which the SH3 is found may therefore be a strong determinant of its ability to bind their recognition motif, as recently shown by Dionne et al. 2021 for more distantly-related SH3s. We therefore performed experiments with the SH3s free of their protein context.

### Free SH3s lose paralog-specific PPIs pattern highlighting the role of protein context on SH3-dependent PPIs

The SH3s were fused directly to the DHFR fragments to measure PPIs, which has been done for instance to identify binding motifs (Wu et al. 2007; Tonikian et al. 2009). All the SH3 variants described previously were expressed by fusing them to the DHFR F[1,2] and we tested PPIs with the preys for which we successfully identified a binding motif (n = 7). The free SH3s were all tagged at either terminus to test the effect of the position of the DHFR fragment on the SH3s PPIs. The SH3 coding sequences were inserted at a genomic locus regulated by the GAL1 promoter and the expression was under the control of a synthetic circuit induced by β-estradiol (Aranda-Díaz et al. 2017) (Figure 5A). This allowed us to test PPIs at various bait expression levels to ensure that we measure PPIs below PCA signal saturation. Analysis of the data shows that the DHFR F[1,2] C-tag empty construct causes less background growth than the DHFR F[1,2] N-tag empty construct (Wilcoxon, p < 2.2×10^-16^), so we focused on the DHFR F[1,2] C-tag constructs.

**Figure 5.**
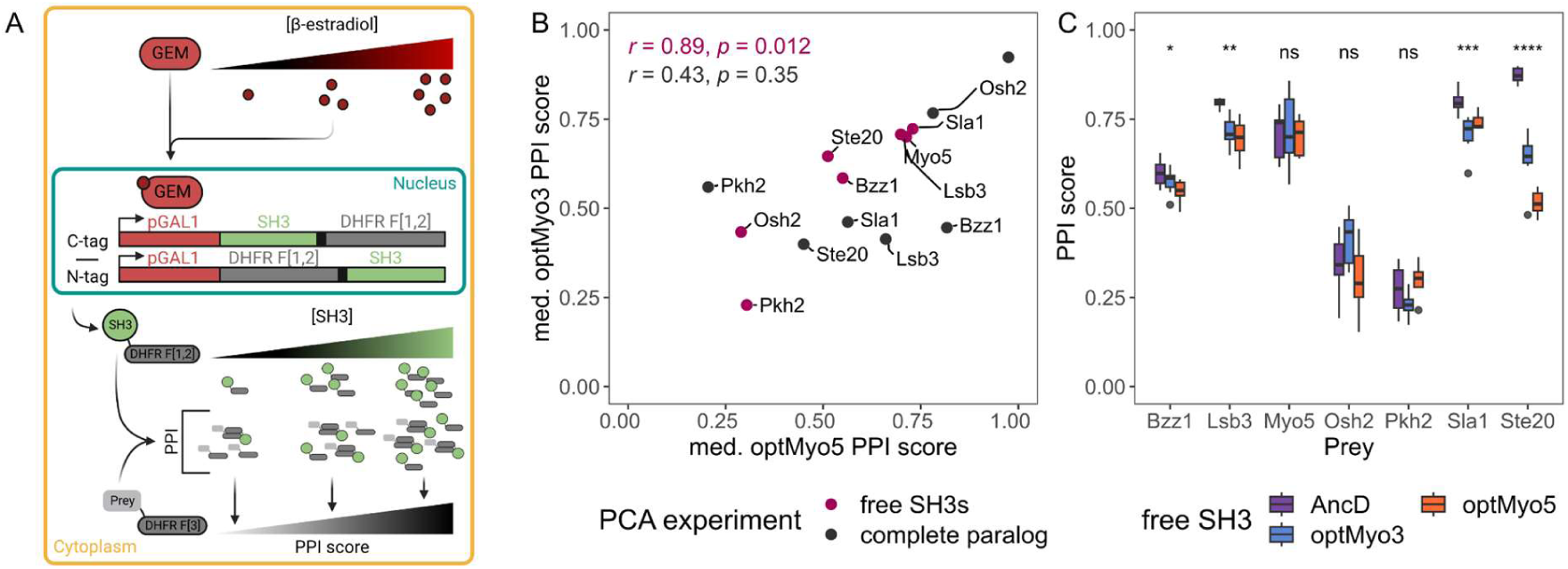
Characterization of free SH3 binding profiles. **A.** Schematic representation of the DHFR PCA experiment using free SH3s with an inducible promoter. The transcription factor GEM binds to β-estradiol and induces the expression of the SH3 coding sequence fused with the DHFR F[1,2] via the *GAL1* promoter. The expression of the construct increases with the amount of β-estradiol, leading to an increase of the DHFR PCA signal. **B**. Scatter plot comparing the PPI profile of the free paralogous SH3s and that of the complete paralogs (entire proteins with the optSH3s). Spearman correlation coefficients (r) and p-values for each PCA experiment are shown. **C**. Boxplot of the PPI scores for several preys (x-axis) with three SH3 variants. Kruskall-Wallis tests compare the scores among the three variants for each prey. P-value significance levels are labeled at the top of each group (ns : p > 0.05, * : p <= 0.05, ** : p <= 0.01, *** : p <= 0.001, **** : p <= 0.0001).

We tested multiple expression levels by changing β-estradiol concentrations and we observed that median PPI scores reach saturation in a prey-dependent manner. To compare the different SH3 variants, we choose a β-estradiol concentration in the range of maximum sensitivity of the assay, below saturation (see methods, Figure S9B). The difference of median PPI score Δ(WT preys - motifΔ preys) in the complete paralogs measured above and the free SH3s shows a significant correlation for optMyo3 (Spearman, r = 0.89, p = 0.012) and the same tendency for optMyo5 (Spearman, r = 0.64, p = 0.14, Figure S9C) suggesting that the free SH3s bind the preys in a similar manner as the SH3s in their host protein context.

The comparison of the median PPI scores between free optMyo3 and free optMyo5 shows a strong significant correlation (Spearman, r = 0.89 p = 0.012, Figure 5B). We observe no bias in interaction strength toward Myo5 SH3 compared to Myo3 SH3 when they are expressed isolated from the proteins (Figure 5B). This supports our results of the swapping experiment that showed that the affinity of the Myo5 SH3 for the prey tested is not higher than the affinity of the Myo3 SH3. However, we tested only seven preys in this experiment, which reduces the resolution to observe divergence. Even if we find that the free paralogous domains generate similar PPIs, the free SH3s do not recreate the paralog SH3-dependent PPI profiles. For example, the PPIs of the paralogs with Osh2 are very strong and are highly affected by the SH3 depletion (Figure 3C), but the free SH3s are not able to generate as strong PPIs (Figure 5B). This can be explained by the competition happening between the free SH3 and the extant paralogs for the binding of their target peptides in those strains.The strains expressing the free SH3s are also expressing the two WT extant paralogs.

As observed previously, the SH3s are causing differences in the PPI scores, but, unlike in the experiments with the entire proteins, we do not detect a systematic decrease in PPI scores of the free AncD compared to the free optSH3s (Figure 5C). In some cases, free AncD creates stronger PPIs than the free optSH3s. Even if it seems that the N-tag free AncD loses PPIs, the statistical test is not significant (Figure S9D, Kruskall-Wallis, p = 0.44), probably because of the limited number of preys included in the experiment and the generally low signal for this construct (Figure S9D). Likewise, there is no apparent decrease in PCA signal for the C-tag free AncD (Figure S9D, Kruskall-Wallis, p = 0.27). This validates the docking results, which predicted that the binding affinity of AncD would be similar to those of the extant domains.

An additional finding we wanted to confirm with these experiments was the stronger interaction between Ste20 and AncD paralog variants when the complete paralogs were considered. We identified the binding motif mediating this PPI (Figure S7C) and we observed a stronger interaction with the free AncD compared to the optSH3s (Wilcoxon, p = 1.6×10^-4^, Figure 5C). However, it is difficult to fully rationalize this result as molecular docking does not predict a higher affinity between AncD and Ste20 motif than with the extant SH3s (Figure 4C). Indeed, it is possible that AncD and Ste20 do not interact as a typical SH3-peptide interaction, thus the docking structure restraints used would not be appropriate to study AncD-Ste20 interaction.

Overall, while ancestral SH3s tend to lose PPIs when introduced in the extant protein, they do maintain an ability to interact with their partners when expressed alone. It is well established that SH3 domains have an intrinsic binding specificity that is encoded in their sequence (Teyra et al. 2017; Brown et al. 2018). However, our results suggest that the ability of the domain to bind motifs on other proteins is also regulated by its host protein via intramolecular interactions or through allostery. Inserting an ancestral sequence into an extant protein could disrupt the regulation mechanism, leading to the general loss of PPIs despite the fact that the domains themselves maintain their ability to bind. This type of regulation implies that coevolution takes place between an SH3 and the rest of its protein such that the intrinsic binding specificity of a domain is not, alone, sufficient to dictate the specificity of the protein.

## Conclusion

We compared the interaction profiles of two recently evolved paralogous proteins, Myo3 and Myo5, that contain SH3s to examine if the sequence divergence of the domains contributed to the divergence in binding to other proteins *in vivo*. Compared to the more than one billion year old origin of the SH3 structural family, these two paralogous domains have recently diverged (Alvarez-Carreño et al. 2021). Our results show that Myo3 and Myo5 have largely overlapping sets of interaction partners but some of their PPIs, which we showed to be SH3-dependent, are biased for one paralog or the other. To test if the SH3 domains are causing these differences in interactions, we swapped the two domains between Myo3 and Myo5. This experiment showed that the two domains are functionally equivalent since they had the same PPI profiles. The equivalent function of the two paralogous SH3s was also confirmed by inserting in the extant paralogs the ancestral SH3 of their last common ancestor. Again, the interaction profiles of the two paralogs were not affected.

To examine further the evolution of the SH3s, we looked at more ancient resurrected domains. We found that in general Myo3 and Myo5 lose their SH3-dependent PPIs when their SH3s were replaced with older ancestral sequences. However, molecular docking showed that the affinity of the most ancient domain reconstructed, AncD, for the proline motifs is not lower than the one of the extant SH3s. Experiments with the SH3 domains isolated from their host proteins revealed similar results. This suggests that although the SH3s may have maintained their binding specificity for millions of years, the way they interact (via intramolecular interactions or through allostery) with their host proteins may have changed. It was previously shown that the modification of a protein domain affects the function of another domain present in the host protein (Cho et al. 2011; Zhang et al. 2023). Thus, allosteric or intramolecular interactions between the host proteins and its SH3 could be disrupted when inserting the ancestral domains into extant proteins, leading to weakened PPIs. Our results thus strengthen the postulate that protein domains are not isolated features of proteins (Melero et al. 2014; Vishwanath et al. 2018) and that evolution is shaped by coevolution between the domains and the rest of the protein.

## Author contributions

PL, DB, AKD and CRL were involved in the design of the study. CRL directed this work. DB performed the phylogenetic analyses and proline motif predictions. PL and AKD performed the experiments. PL performed the molecular docking results and wrote the first draft of the manuscript. DB wrote sections of the manuscript. UD designed part of the molecular biology material. PL, DB, AKD, UD and CRL contributed to revising, reading and approving the final version of the manuscript.

## Data Availability

Strains and plasmids are available upon request. File S1 contains Tables S1-S12. Tables S1-S4 contain (solid and liquid) results from all the DHFR PCA experiments described. Table S5 contains the proline motif prediction information. Table S6 contains the processed flow cytometry data. Table S7 contains a summary of the molecular docking *in silico* experiments. Tables S8 and S9 contain respectively the oligonucleotides and the SH3 DNA sequences. Tables S10 and S11 describe respectively the bait and prey strains used in this study. Table S12 describes all reagents and resources used in this paper. File S2 contains additional details on the methods. File S1-2 and raw data for the DHFR PCA, AlphaFold2 structure predictions, molecular docking, detailed data from the phylogeny, motif orthology analyses and myosins sequence coevolution are available on Dryad at : https://doi.org/10.5061/dryad.sj3tx968m (Data S1-7). Code used to analyze the raw data and to generate all figures can be found at https://github.com/Landrylab/Lemieux_et_al2023.

## Acknowledgments

We thank Soham Dibyachintan, Philippe Després, Romain Durand and Isabelle Gagnon-Arsenault for valuable comments on the manuscript and on the experimental design. We also thank Patrick Lagüe for his insight on molecular docking design. We are also grateful for the support of all the LandryLab members. Some of the figures were created with BioRender and all molecular graphics were performed with UCSF ChimeraX.

## Methods

### Sequence analysis

Multiple sequence alignments (MSA) were performed with MUSCLE v3.8.1551 (Edgar 2004) using the sequences retrieved from the SGD (Cherry et al. 2012).

### Strain construction

Yeast strains used for the DHFR PCA experiments were retrieved from the Yeast Protein Interactome Collection (Tarassov et al. 2008) or constructed. Table S12 (File S1) provides a list of the strain backgrounds and plasmids used to construct the strains. All of the SH3 codon optimized strains (14) were constructed in this study, while WT codon SH3 bait strains (8) were retrieved from Dionne et al. 2021. Bait strains were constructed using strain BY4741 (Mat**a**, *his3Δ leu2Δ met15Δ ura3Δ*) according to Tarassov et al. 2008 method (Table S10 in File S1). Of the prey strains, 6 were individually reconstructed using BY4742 (Matα, *his3Δ leu2Δ lys2Δ ura3Δ*) and 290 were directly retrieved from the Yeast Protein Interactome Collection (Table S11 in File S1). Preys of interest were selected based on a previous study (Dionne et al. 2021) and manually curated from the SGD (Cherry et al. 2012). CRISPR-Cas9 genome editing and homologous recombination were used to construct the paralog variants, the motifΔ preys and the free SH3 constructs. The appropriate media were used to select for the positive clones. NAT (Cedarlane Labs), HYG (Bioshop Canada) and G418 (Bioshop Canada) selections were used at final concentrations of 100 μg/ml, 250 μg/ml and 200 μg/ml respectively. The detailed methods are available in File S2 and illustrated by Figure S10A-C.

### DHFR PCA growth conditions

Bait and prey strains were grown on YPD medium (1% yeast extract, 2% tryptone, 2% glucose, and 2% agar (for solid medium)) containing their specific selection antibiotic. NAT and HYG are used for diploid selection. DHFR PCA selection was made on a synthetic medium (PCA medium, 0.67% yeast nitrogen base without amino acids and without ammonium sulfate, 2% glucose, 2.5% noble agar, drop-out without adenine, methionine, and lysine) containing 200 μg/ml methotrexate (MTX, Bioshop Canada) diluted in dimethyl sulfoxide (DMSO, Bioshop Canada).

### Protein-fragment Complementation Assays and analyses

DHFR PCA protocol is based on Tarassov et al. 2008 and it was published as a visualized protocol (Rochette et al. 2015). The principal purpose of DHFR PCA experiments is to assay the interaction strength between two proteins *in vivo*, by the complementation of the DHFR F[1,2] and DHFR F[3]. The PCA signal is proportional to the amount of reconstituted DHFR in the cell. The detailed protocol is described in File S2. Matting between the baits and preys was performed in random arrays (384 format) on a selective medium resulting in eight or ten replicates for each PPI. The arrays were condensed into 1536 format and then replicated on MTX medium for the assay. Pictures of the microbial growth at the final time point were acquired and used for the analyses.

For each plate picture, the software Pyphe quantified the colonies’ area (‘pyphe-quantify batch --grid auto_1536 --t 1 --d 3 --s 0.05’ (Kamrad et al. 2020)). Raw data from the prey array, diploid selection and PCA selection are available in Data S1. Positions were ignored when there was no growth on the diploid selection or on the prey array. The area values were transformed in log_2_ scale. Then, the background and aberrant data were assessed and removed (3 < log_2_(area) < 13.13, Figure S3A). A standardization using the maximum and minimum logarithm area values on each plate transformed all log_2_(area) on a scale between 0 and 1 (PPI score, see equation below).

If there were two or less biological replicates left after data filtering, the PPI scores were ignored. All the statistical analyses performed on PPI scores are illustrated in the figures or mentioned in the main text. The control group of the statistical tests for the comparison between SH3 variants within one paralog was the optSH3 variant. We choose optSH3 as the control group because ancestral sequences are also codon optimized. Many PPIs were validated in liquid PCA experiments (Figure S3F). The PPI network is visualized with Cytoscape (Figure 1B) (Shannon et al. 2003).

### Comparison of interactome with literature

As a quality control of the data, we compared our dataset with three PPI databases, BioGRID (v4.4.209, (Stark et al. 2006)), STRING (v11.5, (Szklarczyk et al. 2019)) and MINT (Licata et al. 2012). Also, the BioGRID database (v4.4.209 (Stark et al. 2006)) was used as a reference for verification of PPIs detected previously with different experimental methods. The physical PPIs in both orientations (Myo3 and Myo5 as baits and preys) were retrieved and used for comparison (Figure S2A-C).

### Proline motif prediction

Position weight matrix (PWM) specificity models for yeast SH3s were obtained from Tonikian et al. 2009. With the PWMs, a maximum matrix similarity score (MSS) was computed for each SH3-dependent paralog PPI partner (Dionne et al. 2021). The scores are a value between 0 and 1, where 1 represents a perfect match to the PWM (Kel et al. 2003). For each k-mer (k = number of PWM columns) in the preys, MSS were obtained and the maximum MSS k-mer across the whole protein sequence was identified as the predicted binding motif. This procedure was realized for the SH3s of Myo3 and Myo5. The p-value of the prediction was assigned by testing if the maximum MSS score for one prey is higher than the MSSs obtained for 10,000 random peptides. An empirical p-value of 0.05 indicates that the predicted binding motif had a higher MSS than 95% of the random peptides tested against. Proline motif predictions were used to design nine motifΔ preys for which seven were validated experimentally (Figure 1E). The maximum MSS motif positions are shown in Figure S7B (Brennan 2018).

### GFP strain construction and cytometry fluorescence assay

The green fluorescent protein (GFP) DNA sequence and the HPHNT1 resistance cassette were amplified by PCR for genomic integration (File S2). The primers (Table S8 in File S1) were designed to add 40-bp homology arms at each side of the GFP-HPHNT1 amplicon. The DNA fragments were integrated by homologous recombination at the 3’-end of the paralogs replacing the DHFR F[1,2] fragment and the NATMX4 resistance cassette. Hygromycin selection was applied to the cells and the genomic integration was validated by PCR and Sanger sequencing.

Validated GFP-tagged strains were grown at 30॰C in synthetic medium (SC pH 6 HYG, 0.174 % yeast nitrogen base without amino acids and without ammonium sulfate, 2% glucose, 1% succinic acid, 0.6% NaOH, 0.1% MSG) overnight in triplicates. The cultures were diluted in liquid PCA medium without MTX (0.67% yeast nitrogen base without amino acids and without ammonium sulfate, 2% glucose, drop-out without adenine, and 2% DMSO (Bioshop Canada)) to an OD of 0.1, then grown again until the exponential phase was reached (OD = 0.5 - 0.7). The cells were diluted to an OD of 0.05 in sterile water, then the fluorescent intensity of 5000 cells per replicates was measured by a Guava EasyCyte HT cytometer (blue laser, λ = 488 nm).

### Ancestral sequence reconstruction

Ortholog protein sequences of Myo3 and Myo5 were retrieved for fungal species using Ensembl Compara (July 2020, ((Vilella et al. 2009)) (Data S2), ensuring systematic annotations. A MSA of the full-length protein sequences was then produced with MAFFT L-INS-i (v7.453 (Katoh et al. 2005)). Redundant sequences were filtered at a sequence identity threshold of 90% and heavily gapped positions outside of the SH3 were manually trimmed from the alignment. This processed MSA was used to compute a phylogenetic tree using a maximum-likelihood based approach with IQ-TREE2 (Minh et al. 2020) (Figure S1A). The best-fitting amino acid substitution model (LG+G4) was searched automatically within IQ-TREE2 and phylogenetic branch support was assessed using the SH-aLRT test (Anisimova et al. 2011). The tree is represented in Figure S1A (Letunic and Bork 2021). SH3 domain ancestral sequences were reconstructed with the SH3 domain positions from the entire protein’s multiple sequence alignment and the phylogenetic tree using FAST-ML (v3.11, (Pupko et al. 2000; Ashkenazy et al. 2012)). This software has the advantage that it reconstructs insertion/deletions (alongside AA substitutions) in the ancestral SH3 domains. Finally, a recent benchmark of MSA softwares for ancestral sequence reconstruction revealed the algorithm used here (MAFFT L-INS-i) to be one of the best-performing methods for ancestral sequence reconstruction ((Vialle et al. 2018)). The ancestral sequence reconstruction was performed at every tree node using an LG substitution matrix, maximum-likelihood reconstruction of insertions and deletions, and a joint approach for ASR (‘-- jointReconstruction yes’). The nodes of interest were chosen using the taxonomic annotations provided by Ensembl Compara. The ancestral sequences at the node of the most recent ancestor for the clades *Saccharomyces* (AncA_1, AncA_2 and AncA_3, 1^st^ to 3^rd^ most likely sequences), *Saccharomycetaceae* (AncC) and *Saccharomycetales* (AncD) were selected. Some species in the WGD clade lost the additional copy of *MYO3*/*MYO5*, so we also selected the WGD node as another ancestral sequence (AncB). The nodes’ age were estimated with TimeTree (Kumar et al. 2017). The boundaries of the Myo3/Myo5 SH3s were determined from the SMART database (V8.0 (Letunic et al. 2021)). Visualization of the MSAs (Figure S1D-E) were realized with the R package ggmsa (Zhou et al. 2022).

### Molecular docking

The experimental structures of Myo3 (PDB:1RUW) and Myo5 (PDB:1YP5) SH3s were used for the docking. AncD AlphaFold2 (Jumper et al. 2021) best-ranked model was also used (Figure 3C). The structure prediction has a high coverage and pIDDT across the entire structure (Data S3). 28 predicted proline motif structures were computed with AlphaFold2 and the best-ranked model for each motif was used for molecular docking (Data S4) (Tsaban et al. 2022). The molecular docking of the motif predicted structures on the SH3s was done using Haddock2.4 (Dominguez et al. 2003) with parameters optimized for protein-peptide docking (Geng et al. 2017). Ambiguous interaction restraints were defined based on previous work (Janz et al. 2007) (Figure S8A) and all docking parameters are documented in the run.cns files (Data S4). 400 structures resulted from each docking. They were split in clusters according to Haddock2.4 algorithm. The structures were ranked within each cluster, and the clusters were also ranked between each other by the scoring algorithm. The 10 best structures of each cluster, based on Haddock2.4 scoring algorithm, were kept for further computations. The global energy of docking structures was minimized with FoldX (‘RepairPDB’, (Schymkowitz et al. 2005)) 10 times (Usmanova et al. 2018). Then, the interaction energy (ΔG) was computed with FoldX (‘AnalyseComplex’, (Schymkowitz et al. 2005)) for each energy minimized docking structure. The median ΔG across the 10 best structures was computed for each cluster (Figure S8B). All raw and transformed docking data are available in Data S4. Also, all protein visualizations were realized with UCSF ChimeraX (Pettersen et al. 2021).

#### Motif conservation analysis

Orthologs were retrieved from Ensembl Compara (June 2023, (Vilella et al. 2009)) for all 7 preys where the proline motif had been experimentally validated (Figure 1E). We then filtered each set of orthologs to only include species that also contain a Myo3/Myo5 ortholog. Each set of orthologs was aligned using MAFFT L-INS-i, and then the position of the predicted motif in the S. cerevisiae copy was mapped to the multiple sequence alignment. Homologous disordered regions in each ortholog were then retrieved using MobiDB to define the boundaries of the disordered regions (Piovesan et al. 2023). Finally, the best motif in each region was scored using Myo3 and Myo5 PWMs, as described previously (See methods : Proline motif prediction). The Lsb3 predicted motif maps to a structured region, and so a +/-5 alignment window was taken for the motif search range.

**Figure S1.**
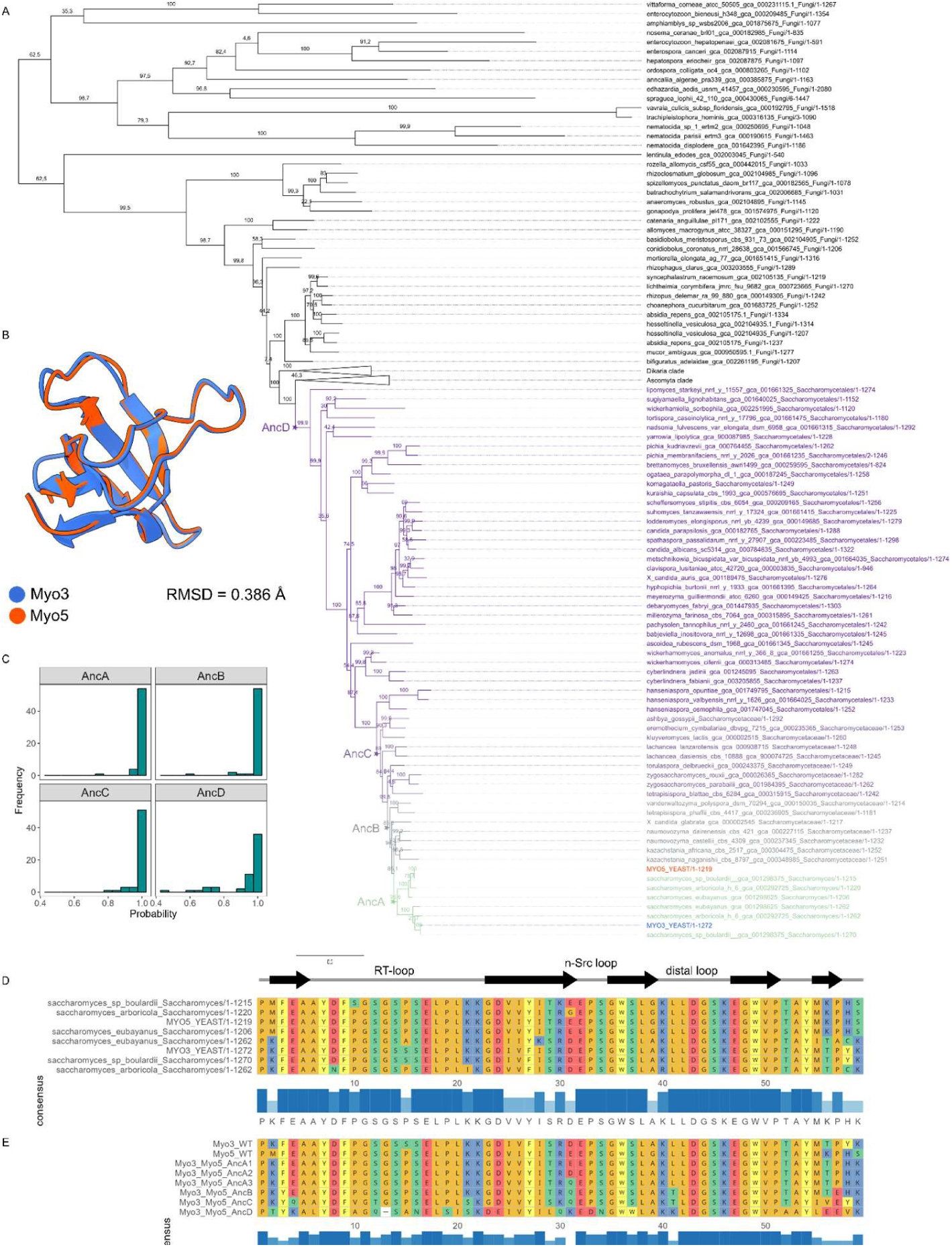
Phylogeny of Myo3/Myo5 orthologs and ancestral sequence reconstruction. **A**. Phylogeny of Myo3 and Myo5 fungal orthologs. The topology obtained respects the fungal taxonomy (Schoch et al. 2020). Myo3 and Myo5 are in mirror clades indicating gene duplication (AncA). The branches used for the ASR are differently colored and the resurrection nodes are labeled by a star (AncA, AncB, AncC, AncD). aLRT values on each branch show overall strong support for the topology. **B.** Superposition of the experimental structures of Myo3 (PDB:1RUW) and Myo5 (PDB:1YP5) extantSH3s shows their high level of similarity (Root Mean Square Deviation of atomic position (RMSD) shown). **C.** Histograms of the highest posterior probability amino acid reconstructed at each position for each ancestral SH3 sequence. The bin width is 0.05. **D.** Multiple sequence alignment of the SH3s belonging to the *Saccharomyces* clade used to reconstruct AncA with conservation chart below. SH3 secondary structures are annotated above. **E.** Multiple sequence alignment of the extant and ancestral reconstructed SH3s. AncA_1, AncA_2 and AncA_3 are the three most likely sequences at the last common ancestor node. AncA_1 is the one used throughout the study (AncA). Position 31 is the only one changing among AncA_1, AncA_2 and AncA_3.

**Figure S2.**
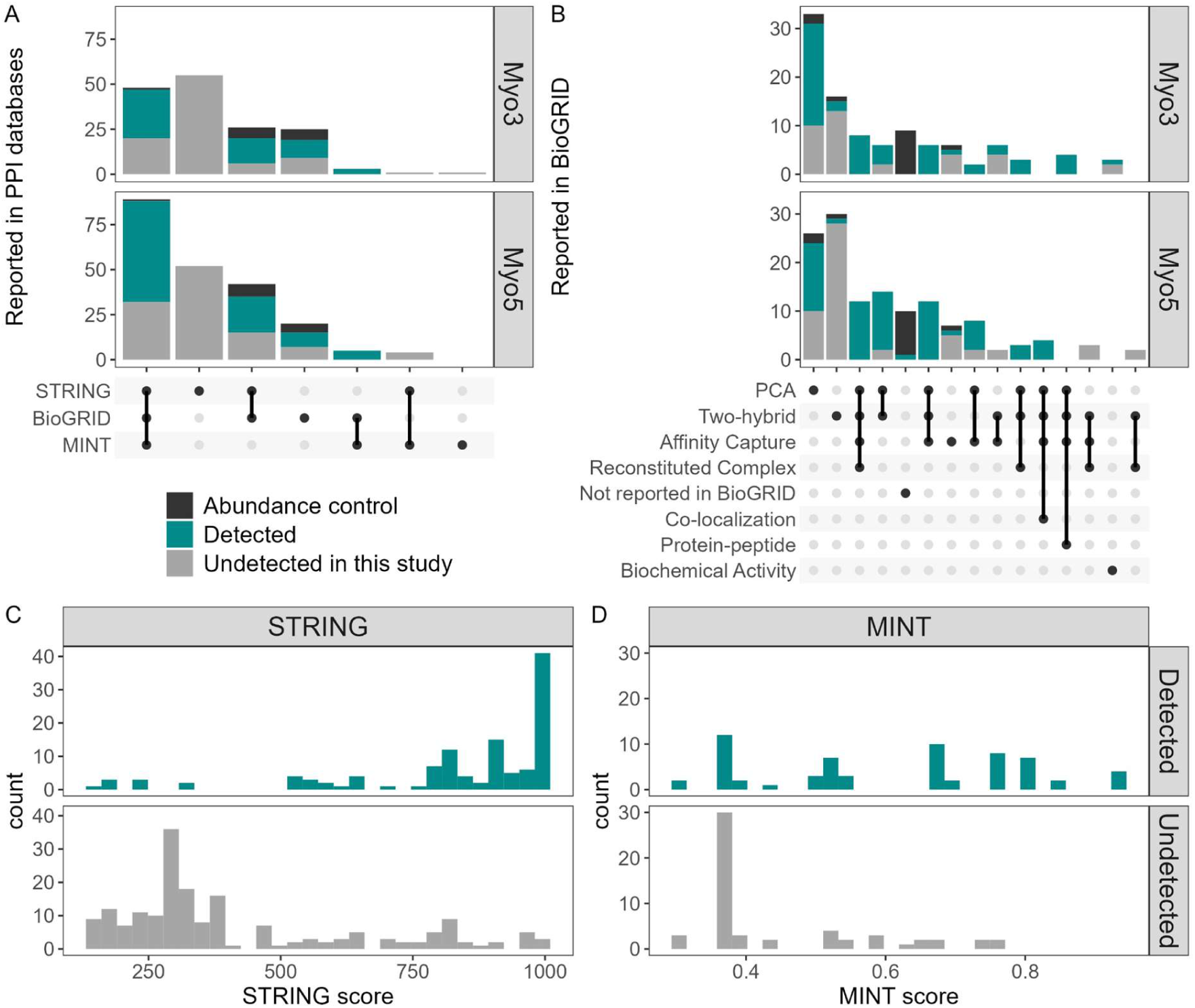
Comparison of the Myo3 and Myo5 PPIs detected in this study with previously reported data. A. Comparison with PPIs deposited in three databases : BioGRID (v4.4.209 (Stark et al. 2006)), STRING (v11.5, (Szklarczyk et al. 2019)) and MINT (Licata et al. 2012). B. Comparison with PPIs deposited in BioGRID for each experimental method reported (v4.4.209 (Stark et al. 2006)). C. STRING confidence score distributions for PPIs detected and not detected in our study. D. MINT confidence score distributions for PPIs detected and not detected in our study.

**Figure S3.**
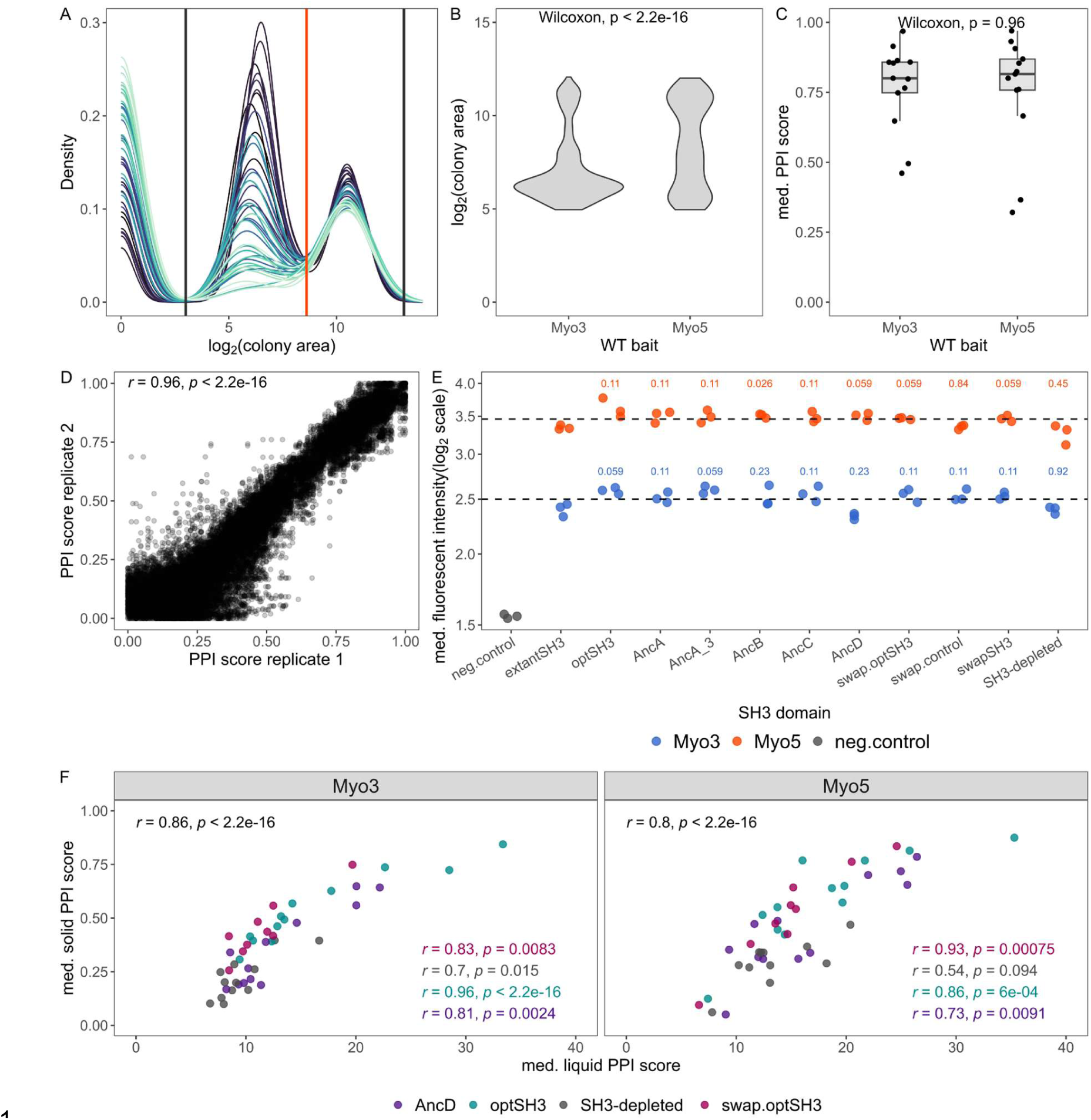
DHFR PCA quality control and validation experiments. **A.** Distribution of the colony areas on a log_2_ scale for the first DHFR PCA experiment (complete paralog with SH3 variants). Each line represents the distribution for one plate of 1,536 colonies. The peak centered at 0 represents the empty positions (no growth) and thus, the background. Colonies with residual growth constitute the middle peak (no PPI) while colonies for which an actual PPI is detected from the peak just above 10. The positions with a colony size between the gray lines are used for normalization (see method). The red line shows the threshold used for identification of true PPIs. **B.** Violin plot showing the distributions of the colony areas on a log_2_ scale for data used for normalization. Myo5 shows larger median colony areas than Myo3 (Wilcoxon, p < 2.2×10^-16^). **C.** Boxplot comparing the median PPI scores of the abundance control with the WT baits. No significant difference is observed (Wilcoxon, p = 0.96). **D.** Scatter plot comparing PPI scores obtained for both technical replicates. D. Protein abundance of the various constructs measured by flow cytometry of GFP fusion proteins. The median of fluorescence intensity for 5000 cells in three replicates is shown. The mean of all SH3 variants for each paralog is represented by a dashed line. Multiple Student’s t-test p-values, Benjamini-Hochberg corrected, comparing the extant paralogs (extantSH3) with the paralogs containing SH3 variants are labeled. F. Scatter plots showing median DHFR PCA PPI score obtained from growth on solid medium plates and in individual liquid cultures from replicated crosses (see method). Growth in liquid cultures was quantified by the logistic area under the curve (AUC) after 65 hours of growth. Different colors used to show the SH3 variants in both paralogous contexts. Specific and global Spearman correlation coefficients (r) are shown. The preys chosen for this experiment are described in Table S4 in FileS1.

**Figure S4.**
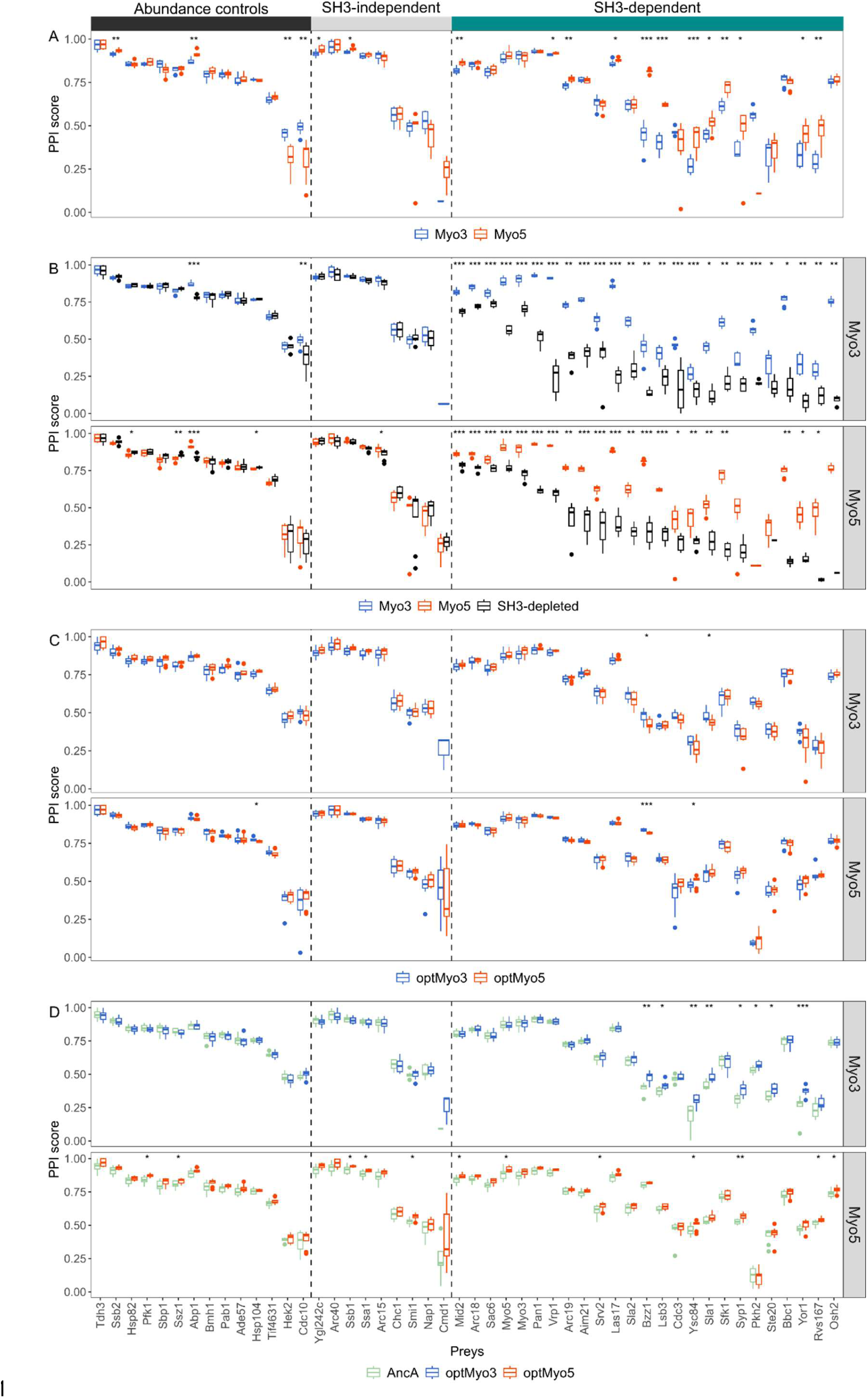
Detailed PPIs comparison between paralogous variants. The colors indicate the SH3 variant for Myo3 paralog (top) and Myo5 (bottom). P-values from Benjamini-Hochberg corrected Wilcoxon tests are shown (* : p <= 0.05, ** : p <= 0.01, *** : p <= 0.001, **** : p <= 0.0001) for each pair. Each interaction was tested in 8 biological replicates. **A.** Extant paralog PPIs comparison. **B.** Identification SH3-dependent interaction partners for each paralog. **C.** SwapSH3 paralog variants PPIs comparison with optSH3 variants**. D.** AncA paralog variants comparison with optSH3 variants.

**Figure S5.**
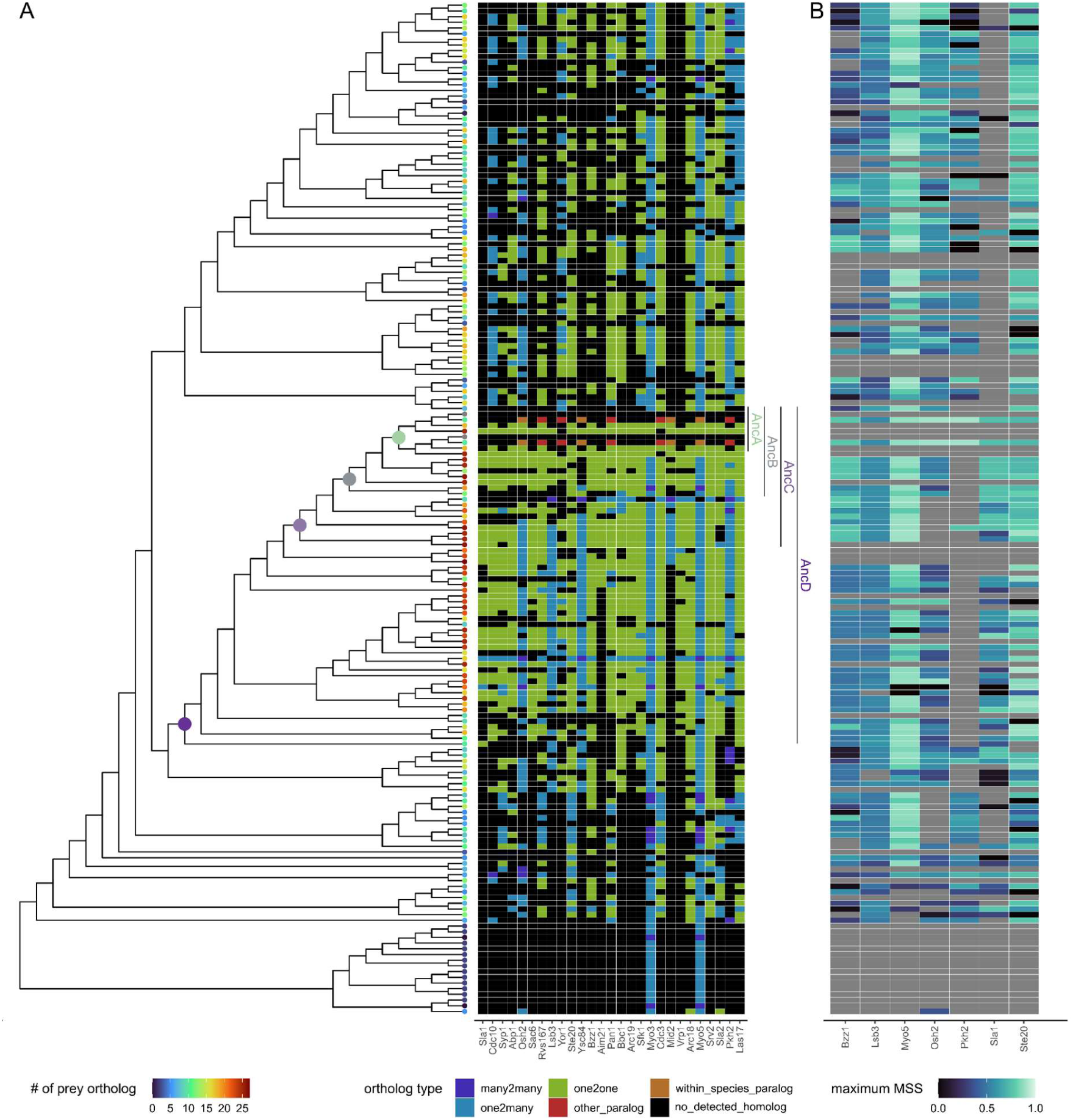
Orthology analysis of Myo3 and Myo5 SH3-dependent interaction partners. **A**. Schematic phylogeny of Myo3 and Myo5 with the tip of its branches colored according to the number of orthologs for the SH3-dependent preys found in each species (left). The orthology information was retrieved from EnsemblCompara (Vilella et al. 2009). The nodes used for the ancestral reconstruction are illustrated with colored circles and the clades are labeled at the right of the phylogeny. The heatmap represents the orthology information for each prey in the species included in the Myo3 and Myo5 phylogeny (right). **B.** Heatmap of the Max MSS score of proline motifs computed with Myo3 SH3 PWM proteins orthologous to the preys aligned with the phylogeny (panel A). The maximum MSS were computed also with Myo5 SH3 PWM which correlated to Myo3 SH3 maximum MSS with a r > 0.99 (Data S5). The maximum MSS motif on each S. cerevisiae preys (x-axis) was validated experimentally (Figure 1E).

**Figure S6.**
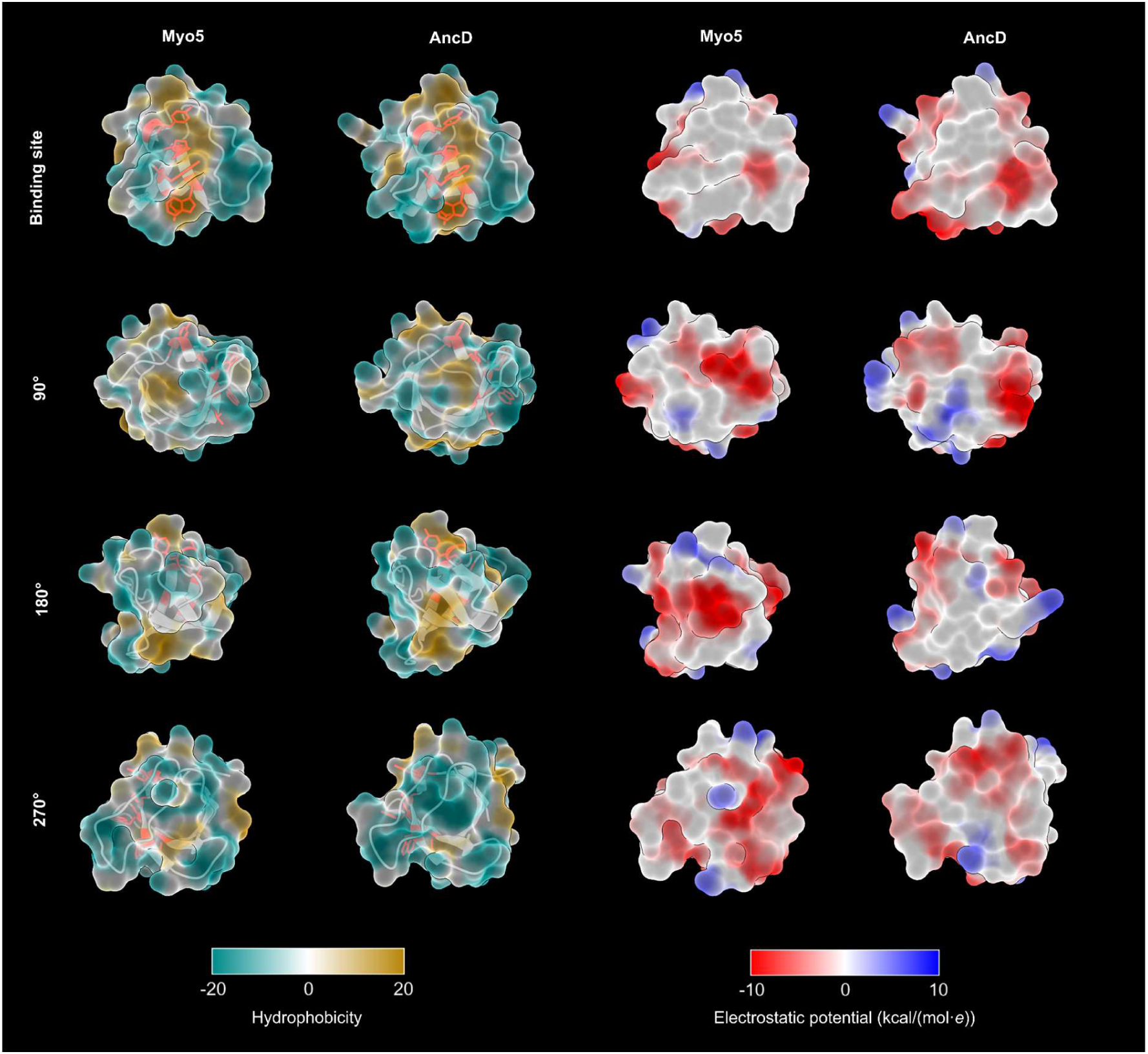
Extant and ancestral SH3 surface properties. Myo5 SH3 and AncD structures viewed in different angles clockwise relative to the Binding site orientation. On the left two columns, hydrophobicity of the surfaces is shown and the secondary structure is displayed under the surface (Laguerre et al. 1997). The residues in red constitute the binding site identified by homology and used to identify ambiguous distance restraints in the molecular docking. The residues of the binding site are identical on AncD and Myo5. On the right two columns, the electrostatic potential of the surfaces is shown. The same analyses were performed using Myo3 SH3 and very similar results to Myo5 SH3 were obtained due to the high sequence similarity between the two extant SH3s.

**Figure S7.**
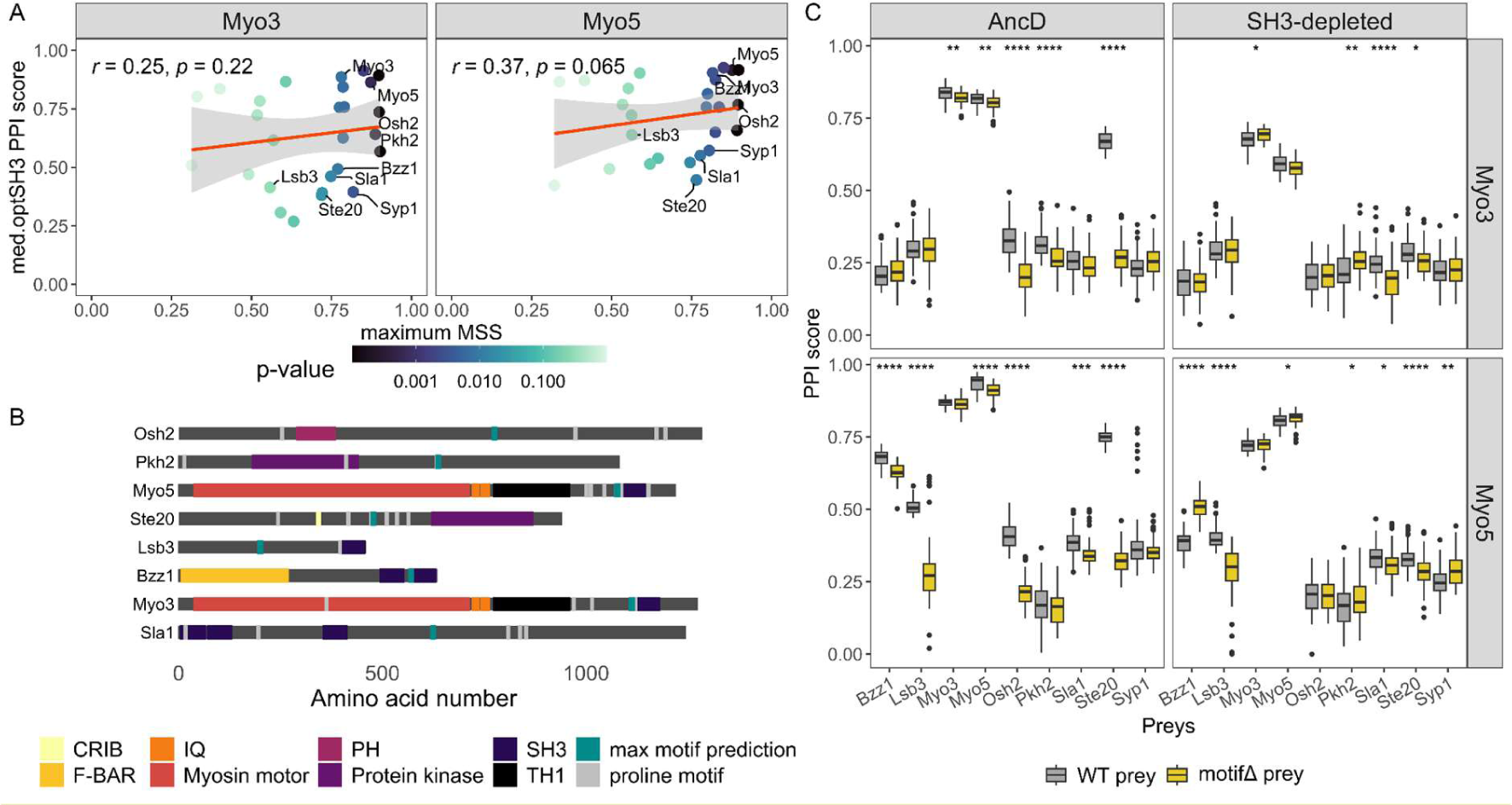
Characterization of the role of proline motif *in vivo*. **A.** Scatter plot comparing the median optSH3 PPI score for each SH3-dependent interaction with its corresponding motif prediction score (maximum MSS). The color scale represents the p-value of the motif predictions which acts as a confidence measure. **B.** Representation of the potential binding motif (PXXP) positions and structural domains on the protein sequence of the tested preys. The motifs with the highest maximum MSS which were deleted in each prey *in vivo* are shown in cyan. **C.** PPI scores of the preys with the binding motifs replaced with a flexible linker compared to the WT preys with paralog variant AncD and SH3-depleted. P-values from Wilcoxon tests are shown (* : p <= 0.05, ** : p <= 0.01, *** : p <= 0.001, **** : p <= 0.0001) for each pair. Each interaction was tested in 48 biological replicates.

**Figure S8.**
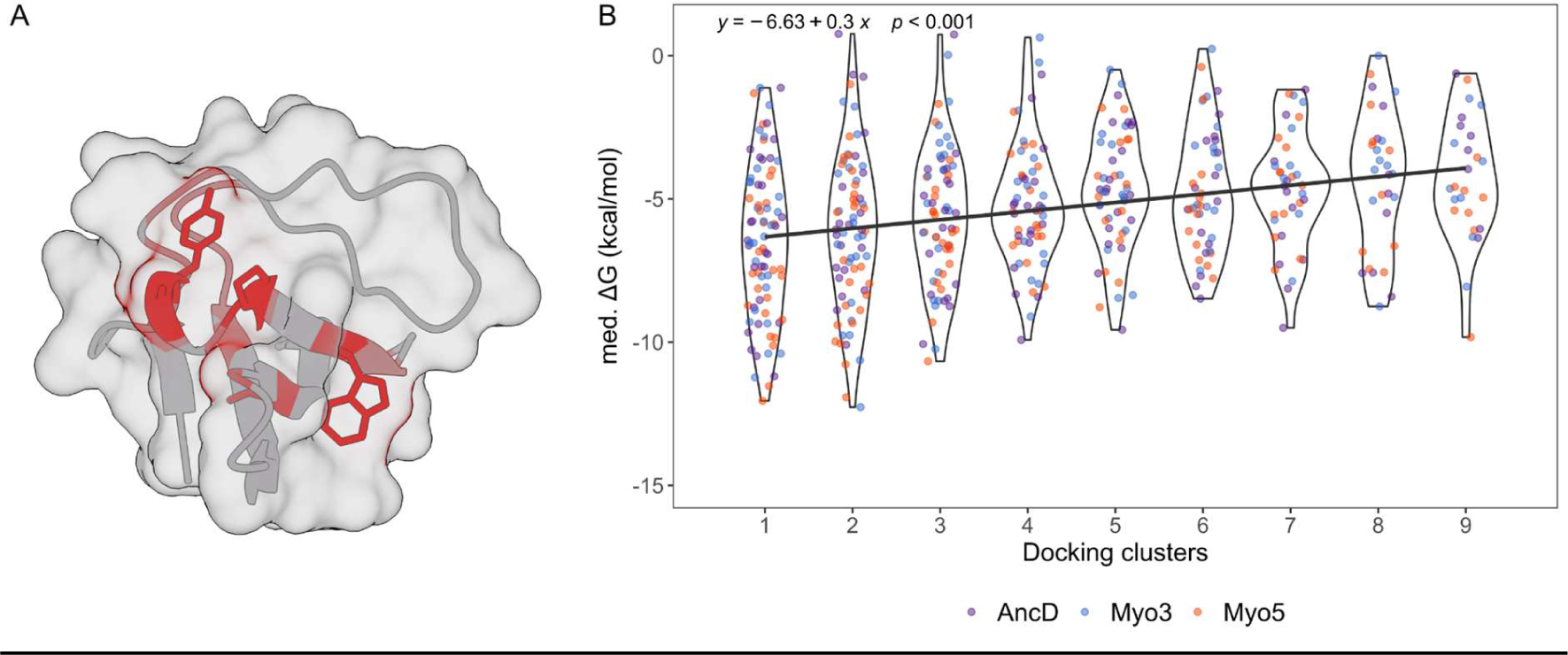
Characterization of the role of the proline motif *in silico*. **A.** Illustration of the ambiguous interaction restraints used for the docking on the Myo3 SH3 structure (PDB:1RUW). Red residues were used as active residues by Haddock2.4. **B.** Violin plot showing the distribution of median ΔGs obtained for the ten best structures of each cluster per peptide (n = 28) for each SH3 variant. The black line is the generalized linear regression of the median ΔGs over each cluster. There is a significant positive relationship between the mean ΔG and the cluster rank for all SH3s tested.

**Figure S9.**
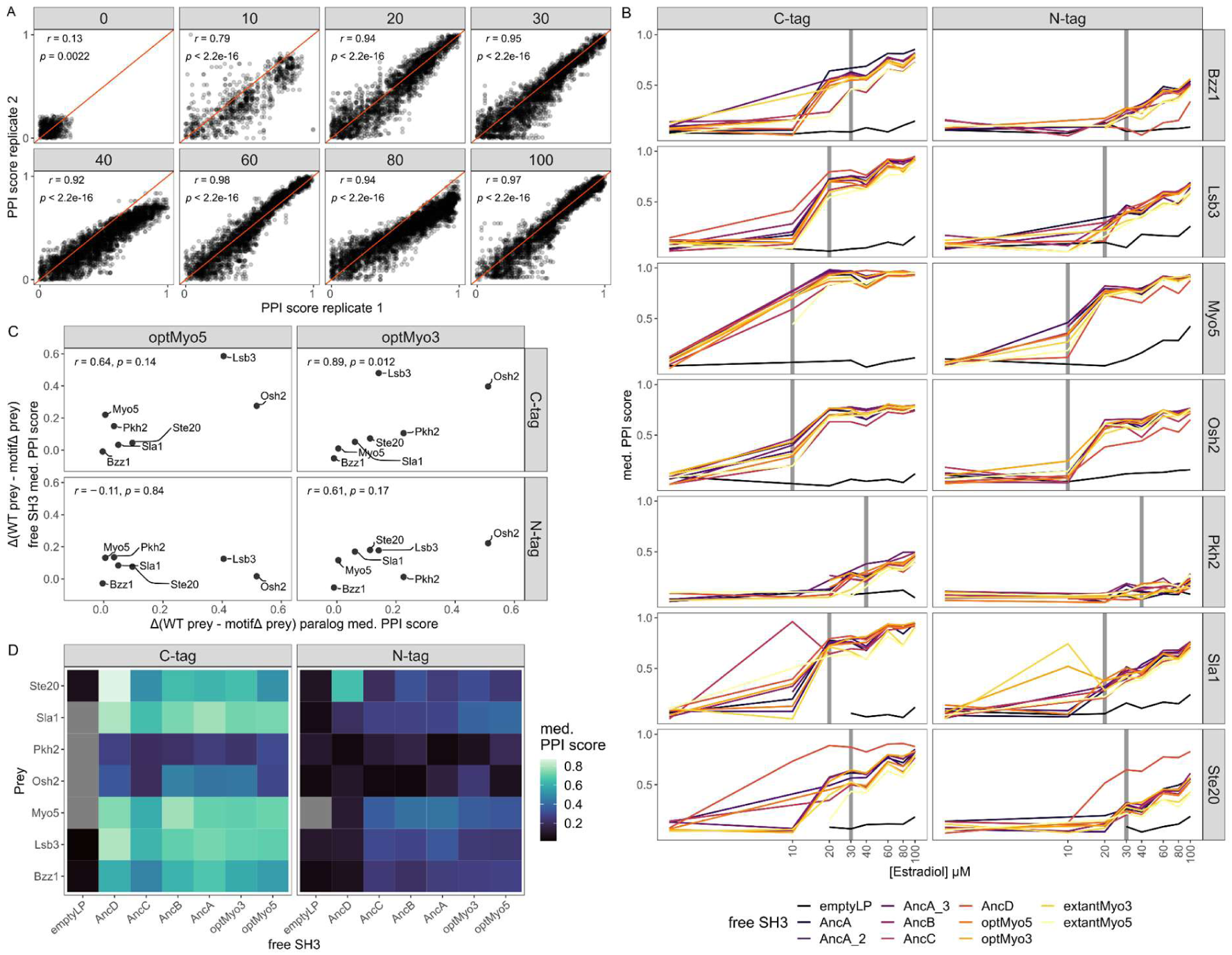
Description of the free SH3s DHFR PCA results. **A.** Scatter plot comparing the replicated PPI scores at each β-estradiol concentration tested (μM). High correlations are observed from 20 μM β-estradiol and higher. For 40 μM and 80 μM conditions, the technical replicates 2 grew less than the replicates 1 which we observe by correlations deviated from the perfect diagonal. Thus, the technical replicates 2 for 40 μM and 80 μM β-estradiol conditions were removed from the analysis. **B.** Plots showing median PPI scores (y-axis) at different β-estradiol concentrations (x-axis, in a log_2_ scale) for each prey. The position of the DHFR F[1,2] fragment is shown independently. The colors represent the different baits. The vertical gray line on each plot shows the β-estradiol concentration at which the sensitivity of the assay was maximal for the prey tested. The median PPI scores obtained at these concentrations are used to compare the different preys. **C.** Scatter plot showing the effect of the motifΔ preys on the PPI scores (Δ(WT prey - motifΔ prey)) for the free SH3s and the complete paralog DHFR PCA experiments. The SH3 variant and the DHFR F[1,2] position are indicated by the row and columns. The prey standard names labeled each data point. Spearman correlation coefficients and p-values are shown for each case. **D.** Heatmap showing the median PPI scores measured between preys (y-axis) and free SH3s (x-axis) tagged with the DHFR F[1,2] in N- or C-terminus and at the β-estradiol concentrations determined in panel B. Disregarding the empty construct, the median PPI scores are not different across the free SH3s (Kruskall-Wallis, C-tag p = 0.27, N-tag p = 0.44).

**Figure S10.**
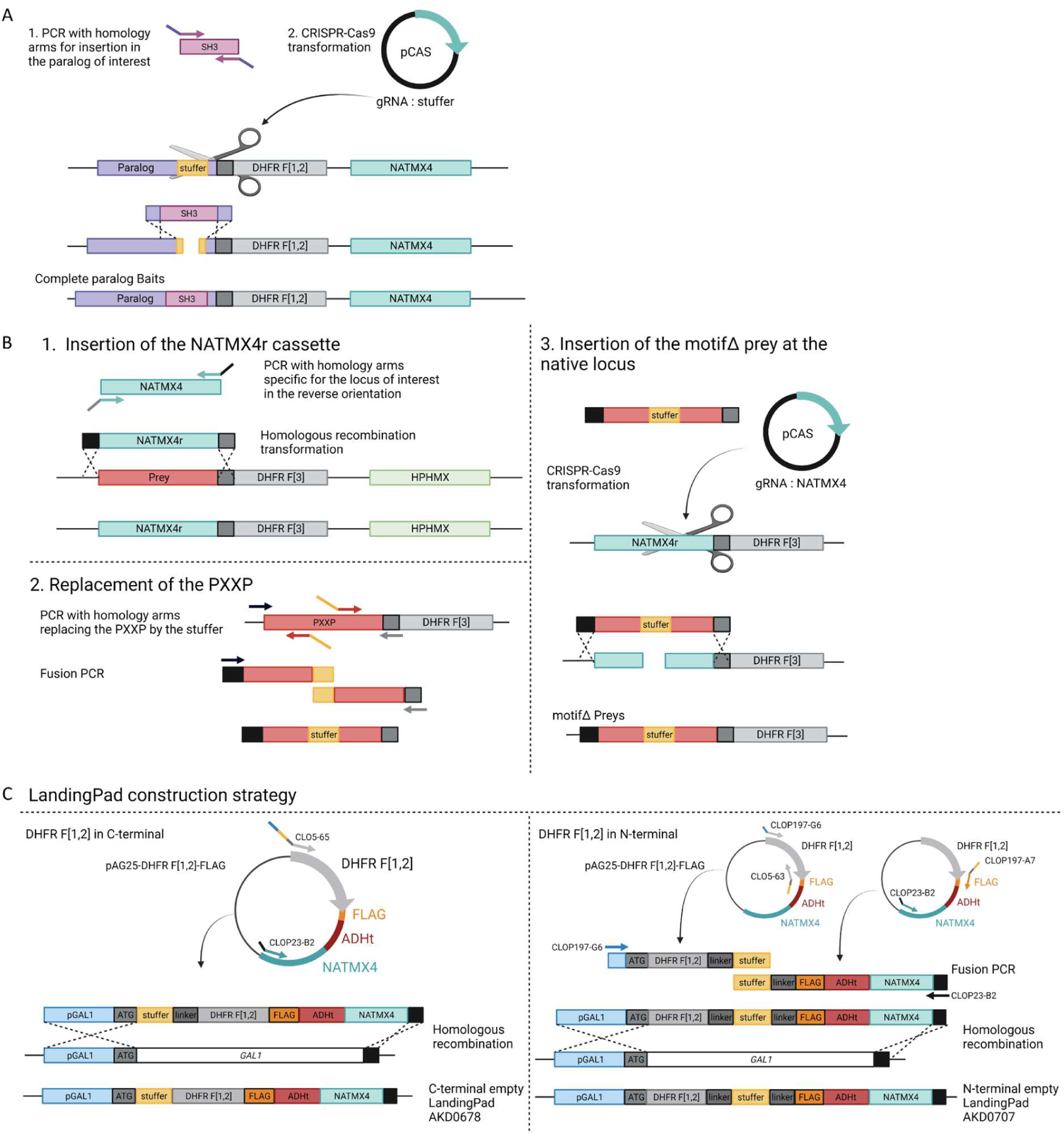
Illustration of the strain construction strategies. **A.** General strategy to insert the SH3 sequences in a genomic locus using a fragment with homology arms (40 bp) for a CRISPR-Cas9 guided homologous recombination (Ryan et al. 2016). **B.** Strategy for the construction of the motifΔ preys. The NATMX4 cassette was amplified for genomic integration at the locus of interest in reverse orientation (NATMX4r). The prey sequence was used for mutagenesis to replace the PXXP motif with a stuffer and homology arms were added to target the locus of interest. This fragment was used for a CRISPR-Cas9 guided recombination using a gRNA targeting the NATMX4r cassette. **C.** Strategy for the construction of the empty LandingPad yeast strains. Fragments were amplified from genomic DNA and from pAG25-DHFR F[1,2] for fusion PCR to add homology arms (40 bp) for integration at the GAL1 locus. Homologous recombination transformations were performed to insert the empty LandingPads in the PL0001 background (Table S12 in File S1). The strategy in panel A is repeated to insert the SH3 sequences at the LandingPad locus.

