## Supplementary material for "Dissection of the role of a SH3 domain in the evolution of binding preference of paralogous proteins": File S2

**File S2 : Detailed methods**

Strain construction

In strains for which the SH3s were replaced by a stuffer DNA sequence (Dionne et al. 2021), CRISPR-Cas9 was used to edit the genomic sequence of the paralogs (Ryan et al. 2016). The stuffer DNA sequence (GGCGGAAGTTCTGGAGGTGGTGGT) codes for a flexible linker (GGSSGGGG). The stuffer design is based on linkers used in structural protein studies and used to join proteins without affecting their structures (Reddy Chichili et al. 2013; Li et al. 2016). PCR was used to create the donor DNA for the CRISPR-Cas9 transformations (pCAS-Stuffer3, gRNA=GGCGGAAGTTCTGGAGGTGG (Dionne et al. 2021)) using codon-optimized SH3 DNA sequences for expression in *S. cerevisiae* (Twist Bioscience) or purified yeast genomic DNA (phenol - chloroform protocol, (Amberg 2005)) as DNA template. The oligonucleotides (Table S8, Eurofins) added 40-bp homology arms encoding for the surrounding regions of the SH3s loci (Figure S6A). All PCRs to amplify DNA for genomic integration were done using the KAPA HiFi HotStart polymerase (Roche) with the following reaction (except mentioned otherwise):

| **Reagent** | **Volume (µL)** |
|---|---|
| Kapa HF buffer 5x | 5 |
| dNTPs 10mM | 0.75 |
| Forward oligo 10µM | 0.75 |
| Reverse oligo 10µM | 0.75 |
| Kapa polymerase | 0.5 |
| Template DNA (15 ng/µL) | 0.75 |
| PCR grade water | 16.5 |
| Total | 25 |

with the following PCR cycle:

<table>
<tr><td>98°C</td><td>5 min</td><td>x1</td></tr>
<tr><td>98°C</td><td>20 s</td><td rowspan="3">x30</td></tr>
<tr><td>60°C</td><td>30 s</td></tr>
<tr><td>72°C</td><td>variable</td></tr>
<tr><td>72°C</td><td>6 min</td><td>x1</td></tr>
</table>

YPD+NAT+G418 (200 µg/mL, Bioshop Canada) selection was applied to cells transformed with pCAS. Colonies were picked randomly to be grown in YPD without selection for plasmid loss. The genomic insertions of the SH3s were confirmed by PCR and the amplicons were sent to sequencing (CHUL Sanger sequencing platform) to confirm the correct genetic modifications. All PCRs to confirm a genetic modification were done using the TAQ DNA polymerase (Bioshop Canada) on purified DNA from picked colonies (Lõoke et al. 2011) using the following reaction:

| Reagent | Volume (µL) |
|---|---|
| TAQ polymerase buffer 10X | 2 |
| dNTPs 10mM | 0.4 |
| Forward oligo 10µM | 0.4 |
| Reverse oligo 10µM | 0.4 |
| $MgCl_2$ 25mM | 1.2 |
| DNA (Quick DNA extraction) | 2 |
| PCR grade water | 13.5 |
| Taq DNA polymerase 5 U/µL | 0.1 |
| Total | 20 |

with the following PCR cycle:

| | | |
|---|---|---|
| 95°C | 5 min | x1 |
| 95°C | 30 s | x35 |
| 54°C | 30 s | |
| 72°C | 1 min | |
| 72°C | 2 min | x1 |

The motifΔ prey strains were constructed using a similar strategy. The predicted binding motif was replaced by the stuffer DNA sequence in the prey strain background, where the DHFR F[3] is fused in C-terminal of the prey. The NATMX4 cassette was amplified in reverse orientation by PCR adding 40pb-homology arms surrounding the loci of interest. Genes coding for the preys were replaced with the NATMX4 cassette by homologous recombination. The Q5 High-Fidelity Polymerase (reaction mix below, New Englands Biolabs) was used for the amplification of the DNA fragments and for the fusion of the motifΔ preys DNA sequences. Then, the genomic NATMX4 cassette was targeted by a CRISPR transformation (pCAS-NAT, gRNA : TTCGTGGTCGTCTCGTACTC) to insert the motifΔ preys DNA sequences at the native prey loci (Figure S6B).

| Reagent | Volume (µL) |
|---|---|
| 5X Q5 Reaction Buffer | 5 |
| dNTPs 10mM | 0.5 |

| | |
|---|---|
| Forward oligo 10µM | 1.25 |
| Reverse oligo 10µM | 1.25 |
| Template DNA | variable |
| Q5 High-Fidelity DNA Polymerase | 0.25 |
| PCR grade water | to 25 |
| Total | 25 |

with the following PCR cycle:

| | | |
|---|---|---|
| 98°C | 30 sec | x1 |
| 98°C | 10 s | x5 |
| 66°C | 15 s | |
| 72°C | 2 min | |
| 94°C | 15 s | x30 |
| 70°C | 15 s | |
| 72°C | 2 min | |
| 72°C | 5 min | x1 |

The strains background for expressing the free SH3s were constructed based on the design of (Aranda-Díaz et al. 2017) creating the PL0001 strain. This strain was again modified to express a genetic construct, designated as a LandingPad (DHFR F[1,2] C-terminal tag : AKD0678 strain, DHFR F[1,2] N-terminal tag: AKD0707 strain) at the *GAL1* locus. The insertion of the SH3 sequences in the LandingPads were completed as described above (Figure S6C). All the strains constructed were confirmed by PCR and Sanger sequencing.

Protein-fragment Complementation Assays and analyses

A pin tool robotic platform (BM5-SC1, S&P Robotics Inc.) was used to perform the PCA screens. From 96-well plates, Matα DHFR F[3] tagged preys were individually arrayed in random positions (384 format) on YPD+HYG solid medium. Then, plates were condensed into 1,536 format on YPD+HYG medium. To remove further border effect, a double border on each edge of the plates is added and used as a control. The control interaction was composed of the bait Lsm8 and the prey Cdc39 (interaction of medium strength). Each prey was present in four or five replicates in the array. Mat**a** DHFR F[1,2] baits were replicated in 1,536 format from a lawn grown on YPD+NAT solid medium. Each bait was crossed twice with the prey array (two technical replicates), resulting in a total of eight to ten biological replicates per PPI. Mating was performed on YPD without selection for 48h at 30°C. Next, two rounds of selection on YPD+HYG+NAT were applied to diploids, each time for 48h at 30°C. As a quality control, pictures of the second diploid selection round were taken after the growth. All images were acquired with a EOS Rebel T5i camera (Canon). The diploids were finally replicated for two rounds on solid PCA selection media. Each time, cells were grown for four days at 30°C in a spImager custom robotic platform (S&P Robotics Inc.). As in previous studies using this method (Dionne et al. 2021), final images from the second selection round were used for image analysis.

A second DHFR PCA experiment was performed using the motifΔ prey strains. The same protocol was followed except for the array design. The extantSH3, optSH3, AncC and SH3-depleted variants in both paralogs (n = 8) were used to create a randomized array in 384 format. The WT prey (n = 9) and motifΔ prey (n = 9) strains were also randomized in a 384 format. The baits and preys were condensed in 1536 format and independently crossed on four plates. The bait and prey arrays were designed to obtain six biological replicates for each PPI per plate (1536 format). After diploid selection, two technical replicates were performed per crossing on PCA medium resulting in 48 biological replicates for each PPI. However, two of the plates were lost due to incubation issues leaving 36 biological replicates per PPI.

A third DHFR PCA experiment was performed using AKD0678, AKD0707, p*GAL1*-SH3-DHFR F[1,2] (n = 10) and p*GAL1*-DHFR F[1,2]-SH3 (n = 10) strains as baits. The same protocol as described above is used, except for the array design. Using the second PCA experiment, 14 preys were chosen for this screen (confirmed proline motif, seven WT and seven motifΔ preys). The mating between the baits and the preys was done in liquid YPD then rearrayed in randomized 384 format. After diploid selection, condensation to 1536 format was performed. There are four biological replicates for each PPI per plate. Successive printing scaled the number of plates to eight. The PCA medium used for this experiment was supplemented with β-estradiol (0 nM, 10 nM, 20 nM, 30nM, 40 nM, 60 nM, 80 nM, 100 nM) (Sigma-Aldrich, #E2758-5G) diluted in ethanol to activate transcription at the genetic constructs at various levels. Each of the diploid plates is used to print two technical replicates on the same PCA condition resulting in eight biological replicates per PPI in each condition.

All pictures were transformed in reverse greyscale so that yeast colonies appear as dark grey (preferred by the image analysis software), then cropped to remove the edge of the omnitrays (ImageMagick Studio LLC). Pyphe (Kamrad et al. 2020), a python toolbox for phenotypic analysis of microbial growth, is used to quantify the colonies' area ('pyphe-quantify batch --grid auto_1536 --t 1 --d 3 --s 0.05') for each plate. Positions that showed no growth on the second diploid selection or on the prey array were removed from the analysis. The area values were transformed in $\log_2$ scale. Then, the background and aberrant data were assessed and removed ($3 < \log_2(\text{area}) < 13.13$ , Figure S2). A standardization using the maximum and minimum logarithm values on each plate ($(\log_2(\text{colony area}) - \log_2(\text{min}))/(\log_2(\text{max})-\log_2(\text{min}))$) transformed all logarithm colony areas on a scale between 0 and 1 (PPI score). PPI scores were considered only if there were two or more biological replicates remaining after data filtering for each PPI.

The analysis of the free SH3 PCA experiment has additional steps. After the processing through Pyphe, the positions that were growing at 0 nM β-estradiol were removed from the dataset at each concentration. We remove those positions because the genetic construct should not enable growth in a β-estradiol depleted media. Also, for the 40 nM and 80 nM β-estradiol conditions, the second technical replicate showed diminished growth (Figure S5), so we removed those data from the analysis. Because all the PPIs are tested at different expression levels of the bait, we had to choose the β-estradiol concentrations for which we considered the PPI scores for each prey. We wanted to choose the expression level of the baits that allows the maximal range of sensitivity of the assay. Thus, we computed the difference in median PPI score between x and x+2 β-estradiol concentration (Δmedian PPI score) associated with the condition x+1. For each PPI, we kept the β-estradiol concentration value x+1 associated with the highest Δmedian PPI score. The β-estradiol concentrations x+1 of each PPI were used to compute the median β-estradiol concentration (rounded up) grouped by prey. This results in one β-estradiol concentration per WT prey at which we considered the PPI scores (Figure S5).

A subset of 71 PPIs detected by DHFR PCA on solid medium were validated using liquid DHFR PCA. The protocol of liquid DHFR PCA was similar to the one in solid. First, the bait and the prey strains were grown respectively in liquid YPD+NAT and in liquid YPD+HYG overnight at 30°C. The mating of the bait and prey strains was realized in liquid YPD without selection (four biological replicates per PPI), then spotted on solid YPD+NAT+HYG (48h at 30°C). Diploids were further grown in liquid SC pH 6 HYG+NAT (0.174 % yeast nitrogen base without amino acids and without ammonium sulfate, 2% glucose, 1% succinic acid, 0.6% NaOH, 0.1% MSG), then diluted to an OD of 0.1 in liquid PCA selection medium using transparent polystyrene 96-well plates (Greiner Bio-One). Growth curves were generated by measuring the OD over four days of incubation at 30°C in an Infinite M Nano plate reader (Tecan). Area under the logistic curve (AUC) was computed with the package R growthcurver (Sprouffske and Wagner 2016) for the first 65 hours of growth. Liquid PPI scores are computed by subtracting the initial OD multiplied by 65 in order to correct the PCA signal for the variations in initial cell inoculate. The Spearman correlation between the results from solid and from liquid DHFR PCA experiments validates the high confidence of the results obtained from growth on solid medium (Figure S2).
